# A *POLR3B*-variant reveals a Pol III transcriptome response dependent on La protein/SSB

**DOI:** 10.1101/2024.02.05.577363

**Authors:** Sandy Mattijssen, Kyra Kerkhofs, Joshi Stephen, Acong Yang, Chen G. Han, Yokoyama Tadafumi, James R. Iben, Saurabh Mishra, Rima M. Sakhawala, Amitabh Ranjan, Mamatha Gowda, William A. Gahl, Shuo Gu, May C. Malicdan, Richard J. Maraia

## Abstract

RNA polymerase III (Pol III, POLR3) synthesizes tRNAs and other small non-coding RNAs. Human *POLR3* pathogenic variants cause a range of developmental disorders, recapitulated in part by mouse models, yet some aspects of POLR3 deficiency have not been explored. We characterized a human *POLR3B*:c.1625A>G;p.(Asn542Ser) disease variant that was found to cause mis-splicing of *POLR3B*. Genome-edited *POLR3B***^1625A>G^** HEK293 cells acquired the mis-splicing with decreases in multiple POLR3 subunits and TFIIIB, although display auto-upregulation of the Pol III termination-reinitiation subunit *POLR3E*. La protein was increased relative to its abundant pre-tRNA ligands which bind via their U(n)U-3’-termini. Assays for cellular transcription revealed greater deficiencies for tRNA genes bearing terminators comprised of 4Ts than of ≥5Ts. La-knockdown decreased Pol III ncRNA expression unlinked to RNA stability. Consistent with these effects, small-RNAseq showed that *POLR3B***^1625A>G^**and patient fibroblasts express more tRNA fragments (tRFs) derived from pre-tRNA 3’-trailers (tRF-1) than from mature-tRFs, and higher levels of multiple miRNAs, relative to control cells. The data indicate that decreased levels of Pol III transcripts can lead to functional excess of La protein which reshapes small ncRNA profiles revealing new depth in the Pol III system. Finally, patient cell RNA analysis uncovered a strategy for tRF-1/tRF-3 as *POLR3*-deficiency biomarkers.

## INTRODUCTION

In eukaryotes, three major RNA types are synthesized by distinct RNA polymerases (Pol): large ribosomal (r)RNA by Pol I, mRNAs and noncoding (nc)RNAs by Pol II, and tRNAs and small ncRNAs by Pol III^1^. With integral initiation and termination subcomplexes that enable recycling, Pol III is the most complex^2–4^. La protein is a ubiquitous component of the Pol III system that binds its nascent transcripts by sequence-specific and length-dependent recognition of their terminal oligo(U) motifs reviewed in ^5–7^. The vertebrate Pol III system has minimalized the length of the oligo(T) DNA transcription terminator to 4Ts in (70% of human) tRNA genes^8–10^. The POLR3E and POLR3K subunits also involved in termination, functionally diverged from yeast^2,11,12^. The type-3 promoter and its trans-acting transcription factors are not found in yeast. In vertebrates, type-3 promoters are limited to few ncRNA genes and require the TFIIIBα subunit BRF2 while type-2 promoters at many more tRNA and ncRNA genes require the TFIIIB**β** subunit BRF1^4,13^.

Some vertebrate Pol III ncRNAs can modulate the expression of specific mRNAs. tRNA fragments (tRFs)^14^, vault (Vt) RNAs and snaRs involve miRNA pathway components not present in yeast^15–20^, and the type-3 U6atac snRNA controls minor-intron gene expression^21^. Primate-specific Pol III transcripts include certain tRNAs as well as snaR-A, VtRNA2-1, and brain cytoplasmic BC200^1,13,22,23^. In sum, human Pol III is more complex and integrally linked to gene expression generally than in yeast at multiple levels^1^.

Most of the >200 pathogenic *POLR3A* and *POLR3B* alleles cause hypomyelinating leukodystrophy (HLD) together with variable impairments in dental and endocrine systems^13,24–28^ see ^29^. Formation of myelin-synthesizing oligodendrocytes are disrupted in mice with POLR3-HLD^30,31^. CNS impairment independent of hypomyelination is common in POLR3-related disorders (-RDs) and can be severe^13,28^. We report homozygous *POLR3B*:c.1625A>G; p.(Asn542Ser) as a mis-splicing variant in a previously undiagnosed patient with neurodevelopmental disease in the absence of hypomyelination, and demonstrate Pol III transcription deficiency in his fibroblasts. CRISPR-editing of HEK293 cells to *POLR3B***^1625A>G^** causes the mis-splicing defect with decreased *POLR3B* mRNA and downregulation of multiple Pol III and TFIIIB**β** subunits, whereas *POLR3E* and La protein are upregulated.

Here, our *in vivo* RNA decay and steady state analyses support the hypothesis that decreased levels of nascent pre-tRNAs in Pol III-deficient *POLR3B***^1625A>G^** cells leads to functional excess of La protein. Assays in *POLR3B***^1625A>G^**cells indicated decreases in pre-tRNA transcription output to only 20-30% of WT cells for tRNA genes with 4T terminators while those with ≥5T terminators were decreased much less and not due to transcript stabilization. Such assays indicate robust decreases in nascent pre-tRNA synthesis after La knockdown (KD) also unlinked to RNA half-life, indicating effects of La on the Pol III transcription process. We show that a known autoregulatory module, in which *POLR3E* mRNA synthesis by Pol II is increased when Pol III transcription of a ncRNA-MIR gene in the *POLR3E* first intron is decreased^32^, is functional in *POLR3B***^1625A>G^** cells and responsive to La KD. Pol III itself mediates transcription interference at *POLR3E* rather than the MIR-ncRNA^32^; hereby validating a role for La in Pol III transcription.

Small-RNAseq showed that *POLR3B***^1625A>G^** and patient fibroblasts exhibit a specific alteration in their profiles of tRNA-derived fragments (tRFs) relative to control cells, that is known of increased La levels^33^. tRF-1s, derived from pre-tRNA 3’-trailers, are specifically increased while tRF-5s and tRF-3s derived from mature-tRNAs are not elevated. These data validate the hypothesis that La protein is in functional excess in *POLR3B***^1625A>G^**relative to control cells and provide evidence of the Pol III transcriptome response in patient fibroblasts. Interestingly but not unexpected, a subset of miRNAs are also specifically elevated.

The Pol III transcriptome response presumably reflects a process that can mitigate impact of *POLR3B* deficiency on cellular function, with notable implications. Presumably because it is associated with coregulation of POLR3 and TFIIIB subunits. The working model is that as Pol III transcripts decreased, La protein resets the ncRNA profile, which in humans also has capacity to reshape the Pol II mRNome. Pol III deficiency consequent to a pathogenic *POLR3B*-variant, upregulates La which contributes to differential expression of tRNA genes. La also impacts tRNA after processing. tRF-1s derived from pre-tRNA 3’-trailers are strikingly increased while the two other major tRFs are not. The results reveal new depth in the Pol III system. Finally, we developed a strategy for tRF pairs to serve as biomarkers for *POLR3*-deficiency.

## RESULTS

A homozygous *POLR3B* c.1625A>G; p.(Asn542Ser) variant was identified in a male proband (P1) evaluated at 22-years of age by the NIH Undiagnosed Diseases Program (UDP)^34,35^. Findings included severe microcephaly and developmental delay; brain imaging at age 16 showed complete myelination, bilateral frontal hypoplasia and thinning of the corpus callosum (**Sup Fig S1A-B**). Sup Table S1 lists additional findings and those of a younger brother, P2, with clinical description in Sup Text 1. Analysis of whole exome sequencing of P1 DNA pointed to *POLR3B* c.1625A>G;p.(Asn542Ser) (NM_018082.6) and *SCRIB* c.890C>G;p.(Thr297Arg) (Sup Tables S2–S3), which were “of uncertain significance” (Sup Table S2)^36^. Although *SCRIB* protein was decreased, actin was not disordered (**Sup Fig S1D-F**) and *Scrib* variants exhibit neural tube defect phenotypes^37,38^, different from P1, P2.

### Analysis of *POLR3B*:c.1625A>G RNA

We confirmed that P1 carries homozygous *POLR3B*:c.1625A>G, for which each parent is heterozygous **(Fig 1A, B)**. Analysis predicted that *POLR3B-*1625A>G SNP may create a new 3’-splice acceptor site near exon-15, prompting inspection of RNA from P1 fibroblasts and two control cell lines (C1, C2). cDNA made with oligo(dT) was followed by PCR with primers in exons 14 and 17. Correctly spliced exons 14-17 would yield a 485 bp product. C1 and C2 produced one band of expected size whereas P1 produced two lower abundance bands **(Fig 1C)**. Sequencing proved that C1 and the P1 upper band represented properly spliced exons 14-17; P1 with 1625G encoding POLR3B-Asn542Ser (**Fig 1B**), and C1 with 1625A, whereas the P1 lower band showed exon-14 spliced to exon-16 with exon-15 precisely skipped (**Fig 1D**, schematically **Fig 1E**). Exon-skipping created a premature stop, *TAG, three codons after the mis-spliced junction (**Fig 1D, E**) that would lead to nonsense-mediated decay^39^.

**Figure 1:**
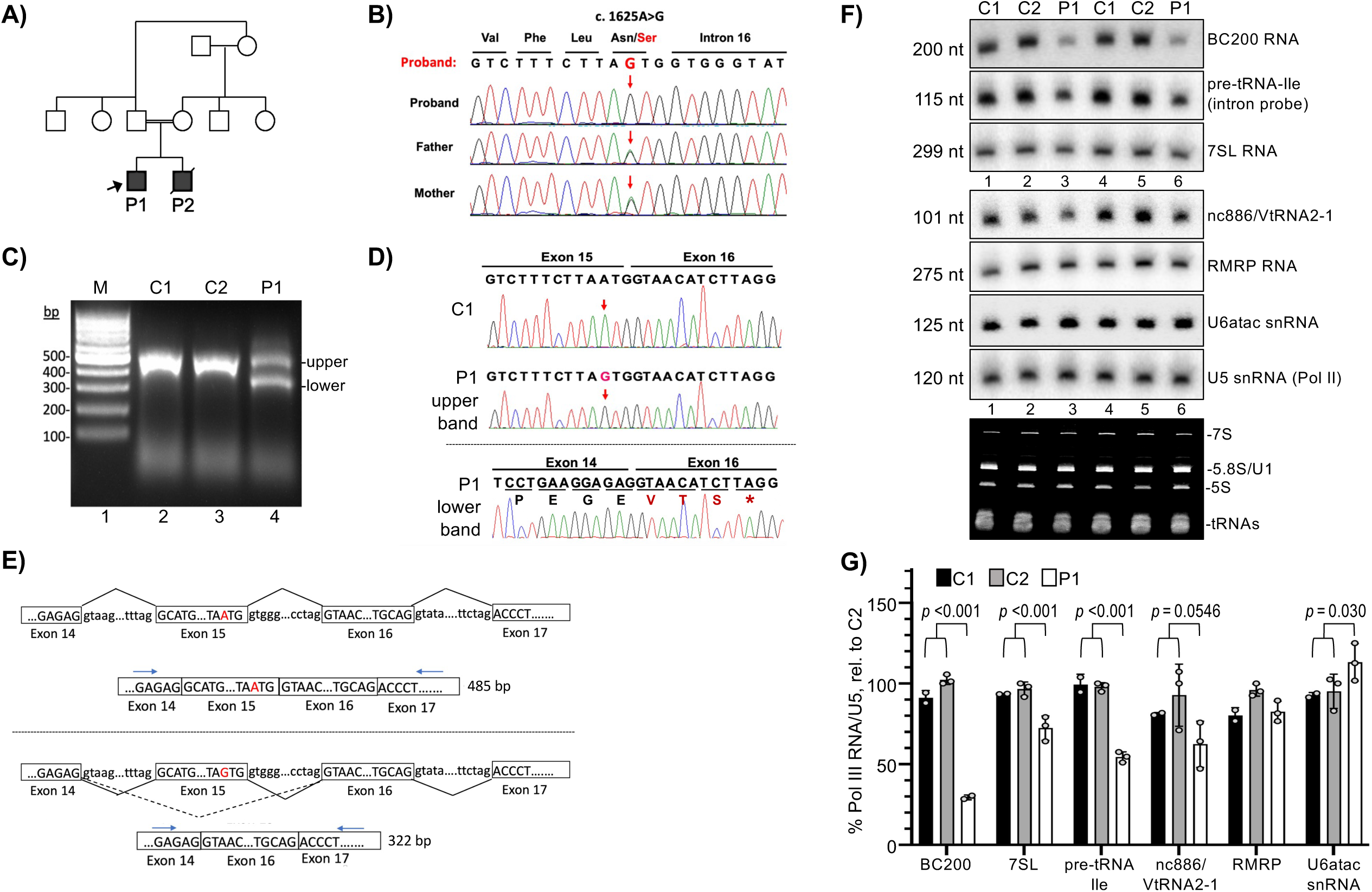
Autosomal recessive *POLR3B*:c.1625A>G; p.Asn542Ser is associated with exon skipping, a premature stop codon, and decreased Pol III transcription products in proband fibroblasts. A) Left, family pedigree; both affected individuals are represented by filled squares; arrow indicates the proband (P1). **B)** Genomic DNA sequencing verifies the homozygous *POLR3B*:c.1625A>G; p.Asn542Ser variant in P1 and that each parent is a heterozygous carrier. Chromatogram shows region around junction of exon-15 and intron-16. **C)** Agarose gel of PCR products from oligo(dT)-primed total RNA isolated from cultured fibroblasts of P1 and 2 control fibroblast lines, C1 and C2. Lane M designates DNA size markers, indicated in base-pairs (bp). The expected size in which exons 14-15-16-17 are correctly spliced is 485 bp. **D)** Sanger sequencing chromatograms of the PCR products obtained from C1 and both products from P1, the upper and lower bands as indicated; downward arrows designate position of the variant nucleotide. The P1 lower band chromatogram includes the one-letter amino acid sequence; the mis-splicing event by this splice-site variant causes precise skipping of exon 15 leading to an in-frame premature stop codon, TAG (asterisk) in exon 16. **E)** Schematics of the introns and exons around c.1625A and 1625G in P1; the schematized products after splicing are also shown. The dashed bent line in the lower panel represents c.1625A>G induced exon 15 skipping. Horizontal arrows indicate PCR primers used. If exon 15 is included, PCR will produce a 485 bp band (upper); if skipped, 322 bp. **F)** Northern blot analysis of total RNA isolated from cultured skin fibroblasts. C1, C2 and P1 above the lanes indicate 2 separate RNA isolations from each, grown in different culture dishes, representing controls from different individuals and P1 cells. A single northern blot was probed for the RNAs indicated to the right. U5 snRNA transcribed by Pol II is a loading control. Bottom panel shows part of the EtBr-stained gel prior to transfer. **G)** Quantitation of replicate northern blot data, normalized by U5 and compared to C2 set to 100%. N = 3 for P1, except for BC200, N = 2. N = 3 for C2, N = 2 for C1. Error bars represent SD. P values were calculated using a two-tailed Student’s two-sample equal variance t-Test; C1 and C2 were considered together and compared to P1.

### P1 cells have reduced levels of Pol III transcription products

Splicing of nascent intron-containing pre-tRNAs occurs early in tRNA maturation with fast turnover. Thus, intron-probed pre-tRNAs serve as a classic assay to assess Pol III transcription in yeast and human cells^40–42^. RNA was probed for pre-tRNA-Ile-TAT1-1, BC200, 7SL, nc886, RMRP, U6atac, and Pol II-transcribed U5 RNA. Pre-tRNA-Ile, BC200 and 7SL were clearly lower in P1 than in C1 and C2 **(Fig 1F-G)**. Because intron-pre-tRNA is an accepted assay for Pol III transcription^41,42^, and given the overlap of signs and symptoms with POLR3-RDs (Sup Table S1), “likely pathogenic” criteria were met for *POLR3B* c.1625A>G; p(Asn542Ser) (Sup Table S2).

### *POLR3B*^1625A>G^ clonal cells

CRISPR-Cas9 genome editing created homozygous *POLR3B*-1625**^A>G^** alleles (hereafter *POLR3B***^1625A>G^**) in two independent clonal HEK293 cell lines named C2 and F4, with the unedited HEK293 parental cells designated wild-type (WT). For *POLR3B* pre-mRNA splicing analysis our HEK293 lab strain was also included; it and WT produced one band of expected size while C2 and F4 produced two (**Fig 2A-B**). The WT and HEK bands, as well as the upper bands from F4 (and C2), represented correctly spliced exons 14-17 with reference nucleotide 1625A (**Fig 2B**). The F4 lower band represented exon-14 mis-spliced to exon-16 with a **\***TAG stop codon (**Fig 2B**, bottom). Thus, C2 and F4 cells exhibited exon-15 skipping as in P1 cells, with POLR3B protein levels at ~25% of WT (**Fig 2C-D**).

**Figure 2:**
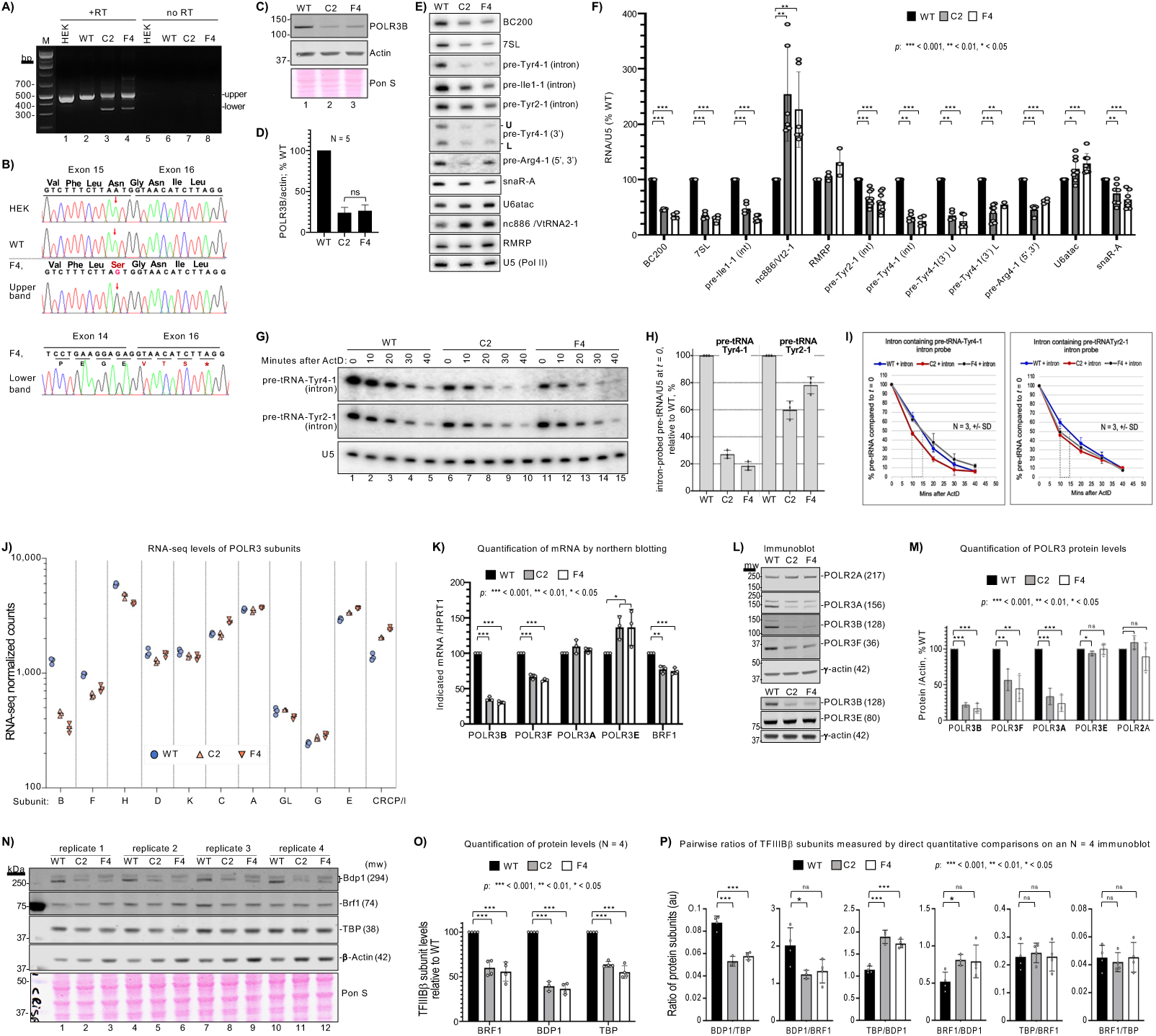
Gene-edited *POLR3B*^1625A>G^ HEK293 cells show exon-skipping, low general Pol III tRNA transcription activity with gene-specific sparing. A) Agarose gel showing PCR products from POLR3B gene-specific oligo-DNA primed total RNA isolated from our lab strain of HEK293 cells (HEK), clonal cell lines C2 and F4, carrying homozygous gene-edited *POLR3B*^1625A>G^, and their HEK293 parent cell line, WT. As in Fig 1C-E, proper splicing leads to a 485 bp RT-PCR product whereas exon-15 skipping leads to a 322 bp product; +RT and “no RT” above the lanes indicates presence or absence of reverse transcriptase. **B)** Sanger sequencing chromatograms of PCR products in A) of HEK, WT, and the upper and lower bands from the F4 sample as indicated. The F4 lower band tracing indicates exon-15 is precisely skipped followed by an in-frame stop codon, TAG. **C)** Immunoblot of WT parent, and C2 and F4 homozygous *POLR3B*^1625A>G^ cell lysates; actin was used as loading control for quantification; positions of MW markers are indicated in kDa. **D)** Quantitation of POLR3B protein levels from 5 separate sample isolations (N = 5) as analyzed in C. Error bars represent the SD, calculated using GraphPad Prism software. **E)** A single northern blot of total RNA isolated from WT, C2 and F4 cells, probed for the transcripts indicated to the right; parentheses, where shown, indicate type of probe used other than to the body, intron, 3’-trailer alone or together with 5’-leader. **F)** Quantitation of northern blot data, normalized by U5. BC200 (N = 4), 7SL (N = 4), pre-tRNA-Ile (N = 6), nc886/VtRNA2-1 (N = 6), RMRP (N = 3), pre-tRNA-TyrGTA2-1 (intron, N = 9), pre-tRNA-TyrGTA-4-1 (intron, N = 5), pre-tRNA-TyrGTA4-1 (3’-trailer U-band and L-band, N = 5), pre-tRNA-ArgTCT4-1 (N = 4), U6atac (N = 8), snaR-A (N = 7). Error bars represent the SD. P values were calculated using two-tailed Student’s two-sample equal variance t-Test. **G)** A single northern blot of an RNA decay time course probed for pre-tRNA-TyrGTA4-1, pre-tRNA-TyrGTA2-1 and U5; harvest time after Actinomycin-D was added to the growth media is indicated above the lanes. **H)** Bar graphs showing steady state, *t* = 0, levels of intron-probed pre-tRNA4-1 (left) and pre-tRNA2-1 (right) in WT, C2 and F4 cells from G) as indicated (N = 3); for each, WT was set to 100%. error bars represent the SD. **I)** RNA decay profiles for pre-tRNA4-1 (left) and pre-tRNA2-1 (right) in WT C2 and F4 cells from G) as indicated (N = 3); vertical dashed-line rectangles extend from the points at which 50% of the RNAs remained (Y-axis) to the time in minutes on the X-axis for estimating half-lives. **J)** Plots of triplicate RNAseq data for the POLR3 subunits along the X-axis. **K)** Quantification of mRNAs analyzed by northern blot in triplicate, using HPRT1 as control; error bars represent the SD. P values calculated using two-tailed Student’s two-sample equal variance t-Test. **L)** Immunoblot analysis; two different blots are depicted and were probed for the proteins indicated to the right (predicted MW in parentheses, in kDa), and each used **γ**-actin as a loading control; <u>mw</u> indicates MW markers on the blot in kDa. **M)** Quantification of the protein levels from immunoblots in L; N = 3; (N = 4 for POLR3E) error bars represent the SD. P values calculated using two-tailed Student’s two-sample equal variance t-Test. **N)** An immunoblot containing four sets of replicate lysates probed by antibodies to proteins listed to the right and visualized using Li-Cor Odyssey infrared imaging system. β-actin was used as a loading control here because γ-actin overlaps with and would obscure quantification of TBP using Abs available for the Li-Cor system. **O)** Quantification of the three TFIIIBβ subunits from immunoblot samples in panel N; N = 4, error bars represent the SD. P values calculated using two-tailed Student’s two-sample equal variance t-Test. **P)** Quantification of the ratios (tabulated independent of actin) of each of the two-way pairs of the TFIIIBβ subunits from the blot in panel N. Error bars represent the SD. P values calculated using two-tailed Student’s two-sample equal variance t-Test.

### Pol III transcriptional deficiency in *POLR3B*^1625A>G^ cells, with tRNA gene-specific effects

Intron-pre-tRNAs, BC200, 7SL, and snaR-A were decreased differentially in *POLR3B***^1625A>G^** vs. WT cells, while the short-lived U6atac and VtRNA2-1/nc886, transcribed from type-3 and a unique Pol III promoter^43–46^, respectively, were increased (**Fig 2E-F**). Intron-pre-Tyr4-1 was decreased approximating POLR3B protein levels in *POLR3B*^1**625A>G**^ cells as were BC200, 7SL and pre-tRNA-Ile1-1. RNAseq revealed misregulation of tRNA-related factors in *POLR3B***^1625A>G^** cells, consistent with Pol III deficiency (**Sup Table S4**). Notably, pre-tRNA-Tyr2-1 and short-lived snaR-A were less decreased than most other ncRNAs in these cells (**Fig 2E, F**). Also, excluding 3’-probed and type-3 RNAs, levels trended lower in F4 than in C2 cells (**Fig 2F**). The tRNA sequences in Tyr2-1 and Tyr4-1 isodecoders are nearly identical, while the leaders, introns, trailers and terminators differ. Intron probing suggests ≥3-fold difference in transcription rate in *POLR3B***^1625A>G^** and WT cells. If the pre-tRNA steady state levels reflect different Pol III transcription, they would be expected to decay/turnover at similar rates in *POLR3B***^1625A>G^** and WT cells. If transcription was the same and posttranscriptional stability accounts for the differences, they would turnover at different rates with ratios of half-lives comparable to the ratios of the steady state levels. We quantified these in triplicate at steady state (*t*=`0)` and during decay after transcription inhibition by actinomycin-D (ActD) (**Fig 2G-I**). We measured 3.4 and 5-fold less pre-tRNA-Tyr4-1 in C2 and F4, respectively relative to WT at steady state (**Fig 2G-H**). The ~0.3 fold-difference in half-life (*t*½, the time when 50% of RNA at *t*=`0` remains) between C2 and WT cannot account for the ~8-fold greater decrease at *t*=`0` **(Fig 2H-I)**; this is also obvious for F4 vs. WT **(Fig 2H-I)**. Thus, Pol III transcription of tRNA-Tyr4-1 is clearly lower in C2 and F4 than in WT cells. By the same measures, pre-tRNA-Tyr2-1 transcription was less decreased in C2 and F4 relative to WT cells than was Tyr4-1, validated by *t*½ **(Fig 2G-I)**. In C2 cells, pre-Tyr2-1 transcription was ≥2-fold higher than pre-Tyr4-1 based on *t*=`0` relative to WT counterparts (**Fig 2H-I)**. In F4 cells, pre-Tyr2-1 transcription was ≥4-fold higher than was pre-Tyr4-1 in F4 relative to WT (based on *t*=`0` relative to WT counterparts) while the *t*½s, both at 10 min, cannot account for this difference (**Fig 2H-I)**. In sum, *POLR3B***^1625A>G^** cells exhibited marked differences in the gene-specific transcription of two tRNA-Tyr genes compared to their transcription in WT cells. While Tyr4-1 transcription is >3 and 5-fold lower in C2 and F4 cells relative to WT, respectively, the Tyr2-1 gene was largely spared from this deficiency in *POLR3B***^1625A>G^**cells.

### Pol III subunits and TFIIIB**β** subunits are downregulated in *POLR3B*^1625A>G^ cells while some are up

RNAseq provided insights into Pol III regulation in *POLR3B***^1625A>G^** cells. Normalized CPMs reflecting Pol III subunit expression showed *POLR3B* was the most decreased in C2 and F4 at ~35% and ~26% of WT, respectively, followed by POLR3F, and POLR3H (Sup Table S5, **Fig 2J)**. Subunits increased in C2 and F4 were CRCP/POLR3I and, less so, POLR3E, whereas POLR3C was increased in F4 only (**Fig 2J**). RNA blotting confirmed subunits B and F decreased, E increased, and confirmed RNAseq data for BRF1 (**Fig 2K**). Immunoblots showed that in contrast to its mRNA, the POLR3A protein was decreased in C2 and F4 to near POLR3B levels, POLR3F was decreased, and POLR3E levels were similar **(Fig 2L-M**).

Side-by-side quantifications of the three TFIIIBβ subunit proteins, BRF1, BDP1 and TBP were performed (**Fig 2N-P**). All three were lower in C2 and F4 vs. WT, relative to β-actin (**Fig 2O**). Direct pairwise ratios confirmed that BRF1 and TBP were comparably decreased in C2 and F4 vs. WT while BDP1 was lower still (**Fig 2P**). Thus, TFIIIBβ subunits are downregulated with POLRs 3B, 3A, 3F and Pol III transcription in both *POLR3B***^1625A>G^** clones. It was unexpected that TBP, which is used by Pols I and II for pre-initiation complex formation, would be decreased in the *POLR3B***^1625A>G^** cells.

### Reduced Pol III activity in *POLR3B*^1625A>G^ cells is rescued by transient increase in POLR3B

Plasmids containing either no insert **(Fig 3A**, lanes 1,3,5), wild-type POLR3B cDNA (2,4,6) or encoding the missense POLR3B-*N542S* encoded by *POLR3B***^1625A>G^** (lanes 7-9), were examined for rescue activity. Immunoblots showing POLR3B and POLR3B-*N542S* expressed at comparable levels suggests similar stability (**Fig 3B**, lanes 1-6). Importantly, neither POLR3B-*N542S* nor POLR3B lowered ncRNAs in WT or *POLR3B***^1625A>G^** cells **(Fig 3A,C**, **S2A,B**), providing evidence of no apparent negative effects. By contrast, raising POLR3B levels increased intron-pre-tRNA levels in C2 and F4 cells relative to no insert **(Fig 3A**, lanes 3-6, **3C**). 7SL was refractory to rescue, BC200 was more responsive in F4 than in C2, whereas levels of short-lived snaR-A were fully reversed in both **(Fig 3A, C)**. Notably, the increase in U6atac in *POLR3B***^1625A>G^** cells relative to WT was reversed after POLR3B expression (**Fig 3C**). VtRNA2-1/nc886 levels were unaffected in WT cells, remained high in C2, and increased in F4 cells with POLR3B expression **(Fig 3C)**.

**Figure 3:**
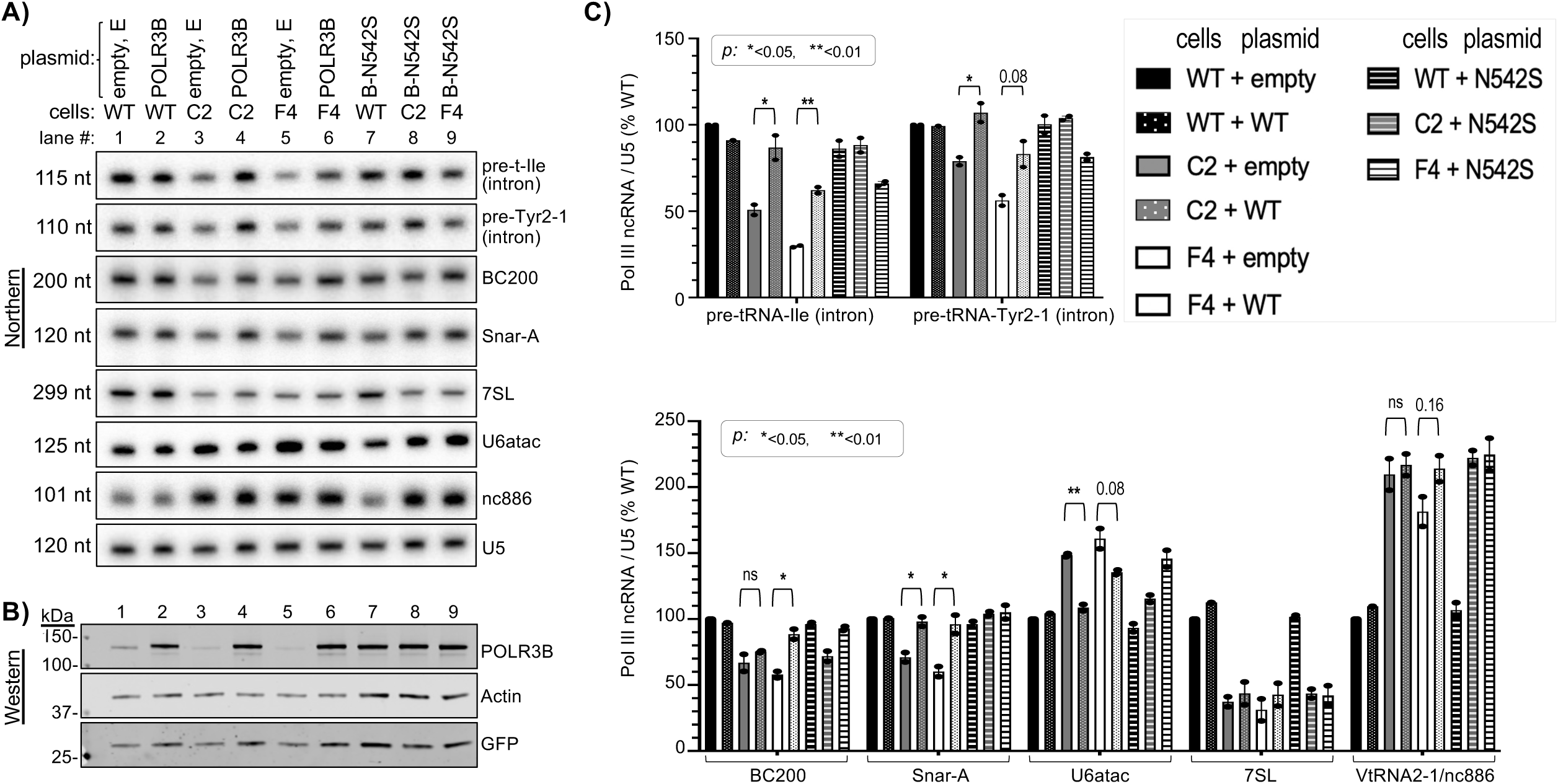
POLR3B expression from plasmid rescues Pol III transcript levels in *POLR3B*^1625A>G^ cells. **A)** A single northern blot of WT, C2 and F4 cell RNAs 48 h after transfection with empty plasmid (E, lanes 1, 3 & 5), POLR3B cDNA (lanes 2, 4 & 6) or POLR3B-*N542S* (B-N542S, lanes 7-9) probed for RNAs indicated to the right. **B)** Immunoblot analysis of proteins from the same cells as in A); GFP was produced from a cotransfected plasmid; Actin is a loading control. **C)** Quantitation of northern blot data, normalized by U5. N = 2, except WT + POLR3B; N = 1. Error bars indicate spread. P values were calculated using a two-tailed Student’s two-sample equal variance t-Test; as indicated, ns = nonsignificant.

POLR3B-*N542S* appeared functionally indistinguishable from POLR3B-WT throughout **(Fig 3A,C)**. This fit with our analysis using a yeast model system^47,48^ in which the homologous Rpc2-N542S substitution did not decrease Pol III activity (**Sup Fig S2C-G**). Thus, the major effect of *POLR3B***^1625A>G^** was mis-splicing with decreased *POLR3B* mRNA, *POLR3B* protein and general Pol III activity.

### Some tRNA genes with **≥**5T vs. 4T terminators are preferentially transcribed in *POLR3B*^1625A>G^ cells

The first tract of ≥4Ts following a tRNA gene sequence is designated the T1 terminator^49,50^. A 4T tract is the shortest functional Pol III T1 terminator^51^ and is found in ~70% of 429 high confidence human tRNA genes^50^. While ≥5T appears to provide only 10-20% higher termination efficiency than 4T^51^, tRNA genes with ≥5T exhibit ≥80% higher Pol III occupancy relative to 4T genes^49^.

Oligo probes specific to introns and 3’-trailers can distinguish nascent pre-tRNAs from various processed species on northern blots to assess their metabolism. Limitations are that only 28 of the 429 genes contain introns of which a fraction can be distinguished from their isodecoders, and of these a fraction have trailers of length and sequence complexity sufficient for detection. We examined two each with ≥5T and 4T in detail, including Tyr4-1 (4T) and Tyr2-1 (5T) (**Fig 4A**, see T**_(N)_**, right column).

**Figure 4:**
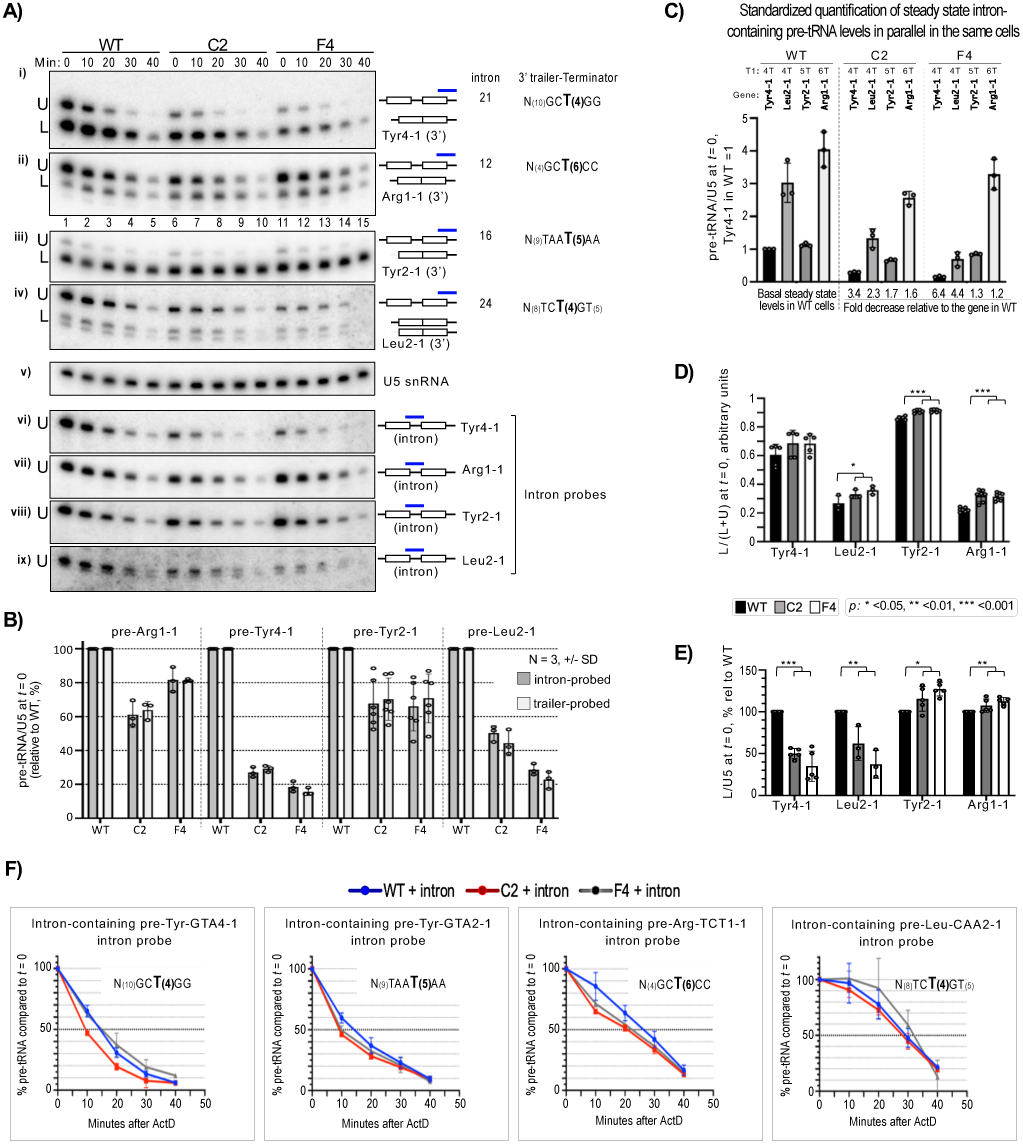
tRNA gene terminator-linked transcription is enhanced in *POLR3B*^1625A>G^ cells. **A)** Northern blot examining transcripts from four tRNA genes using 3’-trailer probes (panels i-iv) followed by intron probes (vi-ix) with U5 RNA as loading control (v). Numbers above lanes indicate time after addition of Actinomycin-D (ActD) to the growth media. Probes are indicated by blue bars over schemas of pre-tRNA intermediates to the right; unfilled rectangles represent exons. The middle column to the right are tRNA gene-specific intron lengths to reflect on mobilities of upper (U) and lower (L) bands (indicated to left of blot panels); the far right column shows gene-specific 3’-trailer features including context and oligo(T) length of the T1 terminator. **B)** Quantitation of the U-bands at *t* = 0 detected by 3’-trailer probes in panels i-iv) and the same RNA species by the intron-probes on the same blots in panels vi-ix); for each tRNA gene, WT was set to 100%. Error bars represent the SD. **C)** Parallel quantification of intron-containing, trailer-probed, pre-tRNAs from steady state, *t* = 0 three replicate experiment samples (N =3); the Tyr4-1 WT was set to 100%. **D)** Quantitation of 3’-trailer-containing spliced pre-tRNA, the L-bands at *t* = 0, in WT, C2 and F4 cells relative to WT, represented by L/(L+U). **E)** Quantitation of the total L-bands at *t* = 0. For D-E, *p* values were calculated using a two-tailed Student’s two-sample equal variance t-Test; represented by asterisks as below panel D; error bars represent the SD. **F)** Pre-tRNA decay profiles derived from quantification of triplicate ActD decay experiments as in A for each of the intron-probed pre-tRNAs indicated; error bars represent the SD. Insets show the gene-specific 3’-trailer/terminator features as in A.

The major difference between the upper (U) and lower (L) bands detected by trailer probes is absence of an intron in L (**Fig 4A**; panels i-iv and vi-ix). Sequential probings and quantifications indicate that the major bands detected by intron probes were indistinguishable from U-bands by trailer probes **(Fig 4A-B)**.

tRNA gene-specific differences between WT vs. C2 and F4 are apparent in **Fig 4A**, i-ii. The U-band nascent pre-tRNA-Tyr4-1 is more abundant in WT vs. C2 and F4, reflective of their relative POLR3B levels. This pattern was not apparent for the U-band nascent pre-tRNA-Arg1-1 in the same cells, which was relatively more abundant in C2 and F4 compared to Tyr4-1 (**Fig 4A**, i-ii). This differential between Tyr4-1 and Arg1-1 in WT vs. C2 and F4 was also observed for intron-probed pre-tRNAs (**Fig 4A**, vi-vii).

Pre-tRNA-Tyr2-1 and pre-tRNA-Leu2-1 exhibit the same major difference as observed for Tyr4-1 and Arg1-1. The U-band pre-Leu2-1 was more abundant in WT vs. C2 and F4, while the U-band pre-Tyr2-1 did not follow this pattern, confirmed by intron probing (**Fig 4A**, iii-iv, viii-ix). In sum, quantification of nascent intron-pre-tRNAs suggests higher Pol III transcription for tRNA-Tyr2-1 and tRNA-Arg1-1 genes than for tRNA-Tyr4-1 and tRNA-Leu2-1 in *POLR3B***^1625A>G^** cells compared to WT probed for introns or 3’-trailers **(Fig 4A-B)**, extending **Fig 2G-H** data.

Quantifications thus far were from each tRNA gene in each cell line, setting WT to 100%. To estimate relative differences for these tRNA genes in the *same* cells we performed standardized quantifications in parallel. This produced measures for each tRNA gene in WT cells which we will refer to as their relative basal activities (**Fig 4C**, left four lanes). This analysis showed that their differential activities in *POLR3B***^1625A>G^** cells were principally unlinked from the relative levels of their basal activities in WT cells. Output from the 4T terminator genes was decreased more than output from the ≥5T terminator genes in the C2 and F4 cells apparently independent of their relative basal activities in WT cells (**Fig 4C**, quantification summarized under lanes). In sum, tRNA gene activities with 4T terminators were accordant with *POLR3B* deficiency, while gene activities with ≥5T terminators were substantially less susceptible to the deficiency. We note that while the differential tRNA gene activities appear linked to their gene-specific properties, these results do not demonstrate that the terminators per se are responsible.

### Evidence of enhanced La activity in *POLR3B*^1625A>G^ cells

La protein binds nascent pre-tRNAs and other Pol III transcripts at their 3’oligo(U) termini and promotes their maturation^52,53,**see**^ ^6,54^. As shown in a later section, La is upregulated in *POLR3B***^1625A>G^** relative to WT cells at RNA and protein levels. Thus, relative to Pol III transcripts, La would be in excess of its ligands in *POLR3B***^1625A>G^** cells.

*POLR3B***^1625A>G^** cells accumulated more intron-less species, L-band, as a fraction of transcript output, L/(L+U), than WT cells for all four pre-tRNAs (**Fig 4D)**. Most distinctively, accumulation of L independent of total was increased in *POLR3B***^1625A>G^**for the ≥5T genes, but decreased for the 4T genes **(Fig 4E),** and not accountable by cell-specific differences in 3’-trailer turnover (**Sup Fig S3C**). These data suggest involvement of La, which binds UU(n)U in a length-dependent manner, following oligo(T) length, though usually shorter^55,56^ see ^7^. Further, nascent-pre-tRNAs were generally processed faster in *POLR3B***^1625A>G^** cells than in WT (**Fig 4A,F**, **Fig S3A,C**), consistent with La protein as a molecular chaperone^52,57^ see ^5^.

### VtRNA2-1 transcription is upregulated in *POLR3B*^1625A>G^ cells

Direct comparison showed increased VtRNA 2-1 to 1-2 in *POLR3B***^1625A>G^**vs. WT cells (**Fig 5A**). We compared VtRNAs at steady state, *t*=`0`, and during turnover (**Fig 5B-E**). Unlike Vt2-1, Vt1-1,2,3 levels were not much altered in *POLR3B***^1625A>G^** cells (**Fig 5C**). Vt2-1/nc886 displayed *t*½ of <90 minutes as published (**Fig 5D**)^45,58^. Small differences in turnover indicate that increased transcription accounts for the ~3-fold higher levels in *POLR3B***^1625A>G^** cells.

**Figure 5:**
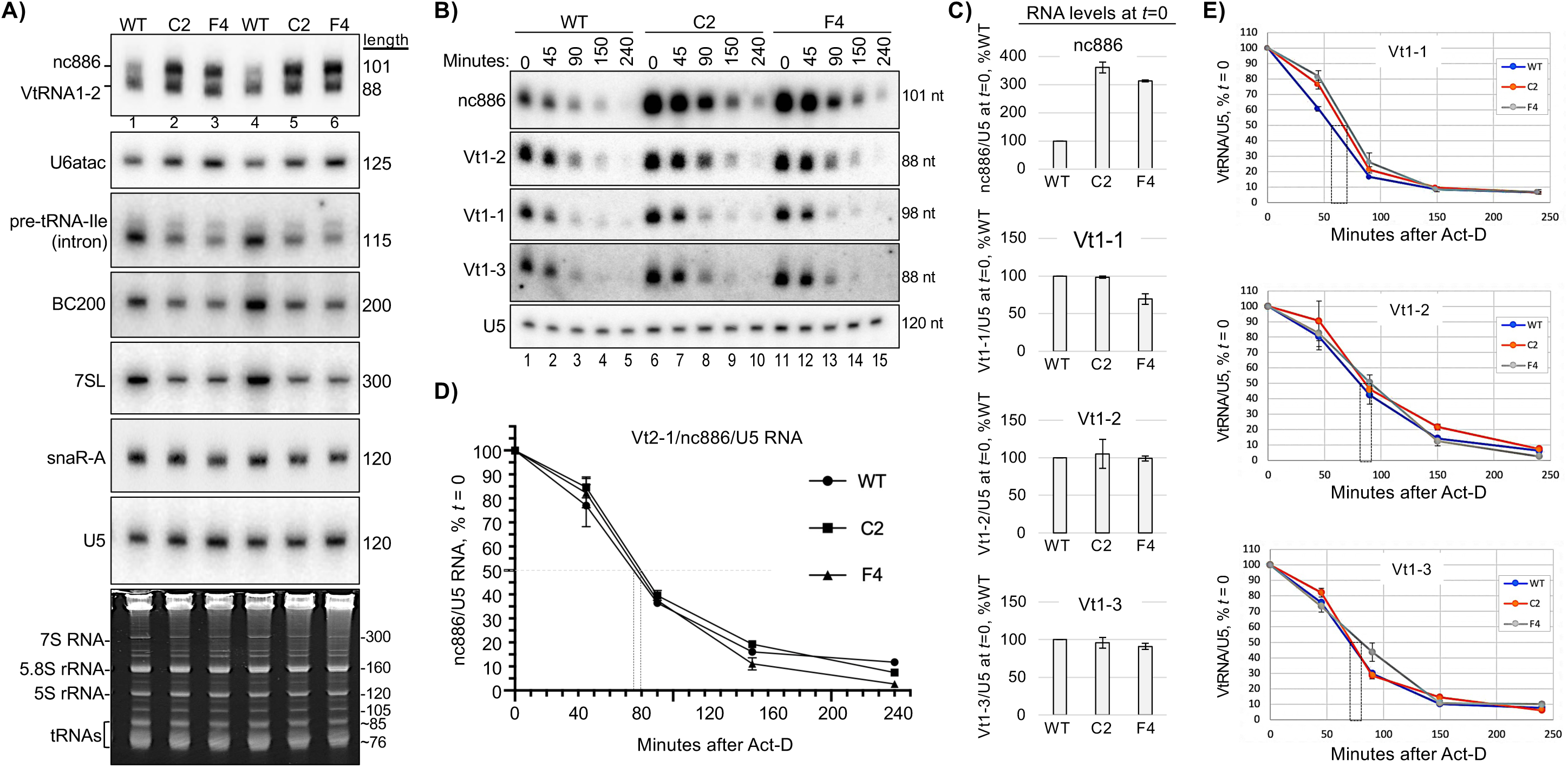
Transcriptional upregulation of nc886 RNA in *POLR3B*^1625A>G^ cells. **A)** Northern blot probed for transcripts indicated to the left. **B)** Time course of posttranscriptional RNA decay in WT, C2 and F4 cells as indicated above lanes with times after ActD addition to the growth media. **C)** Quantification of duplicate data for the ncRNAs in B at *t* = 0 (steady state); error bars indicate spread. **D)** Graphic quantification of duplicate data for nc886 RNA decay; error bars indicate spread and the vertical rectangle extends from the point at which 50% of the RNAs remained (Y-axis), to the time on the X-axis, for estimating half-life. **E)** Quantification of decay data for VtRNAs 1-1, 1-2 and 1-3 as in D.

### La knockdown (KD) decreases cellular Pol III transcription

Two siRNAs against La were compared to negative control (NC) siRNA (**Fig 6A**). La was increased at the protein and RNA levels in C2 and F4 vs. WT siNC cells and decreased in siLa cells (**Fig 6B** upper and lower panels). La-KD effects on Pol III ncRNAs were reproducible in multiple experiments by blotting and quantification (**Fig 6C-E**). La-KD decreased the ncRNAs in **Fig 6D** by 30-65% relative to siNC, generally more by siLa1 than siLa2. VtRNA 2-1, 1-1, and U6atac were decreased more in *POLR3B***^1625A>G^** than in WT cells, while snaR-A was not. The pre-tRNAs were decreased less by La-KD than the ncRNAs in Fig 6D (**Fig 6E**). These ncRNAs were largely spared from the Pol III deficiency in *POLR3B***^1625A>G^** cells or were expressed at levels higher than in WT cells (**Fig 2F**, **Fig 5C**). Tyr2-1 and Arg1-1 were decreased by 20-40% and more in *POLR3B***^1625A>G^** than in WT cells **(Fig 6C, E)**. Pre-tRNA Leu2-1 was not decreased as much as Tyr2-1 and Arg1-1 by siLa1 and even less by siLa2. Tyr4-1 levels were the overall least decreased by siLa.

**Figure 6:**
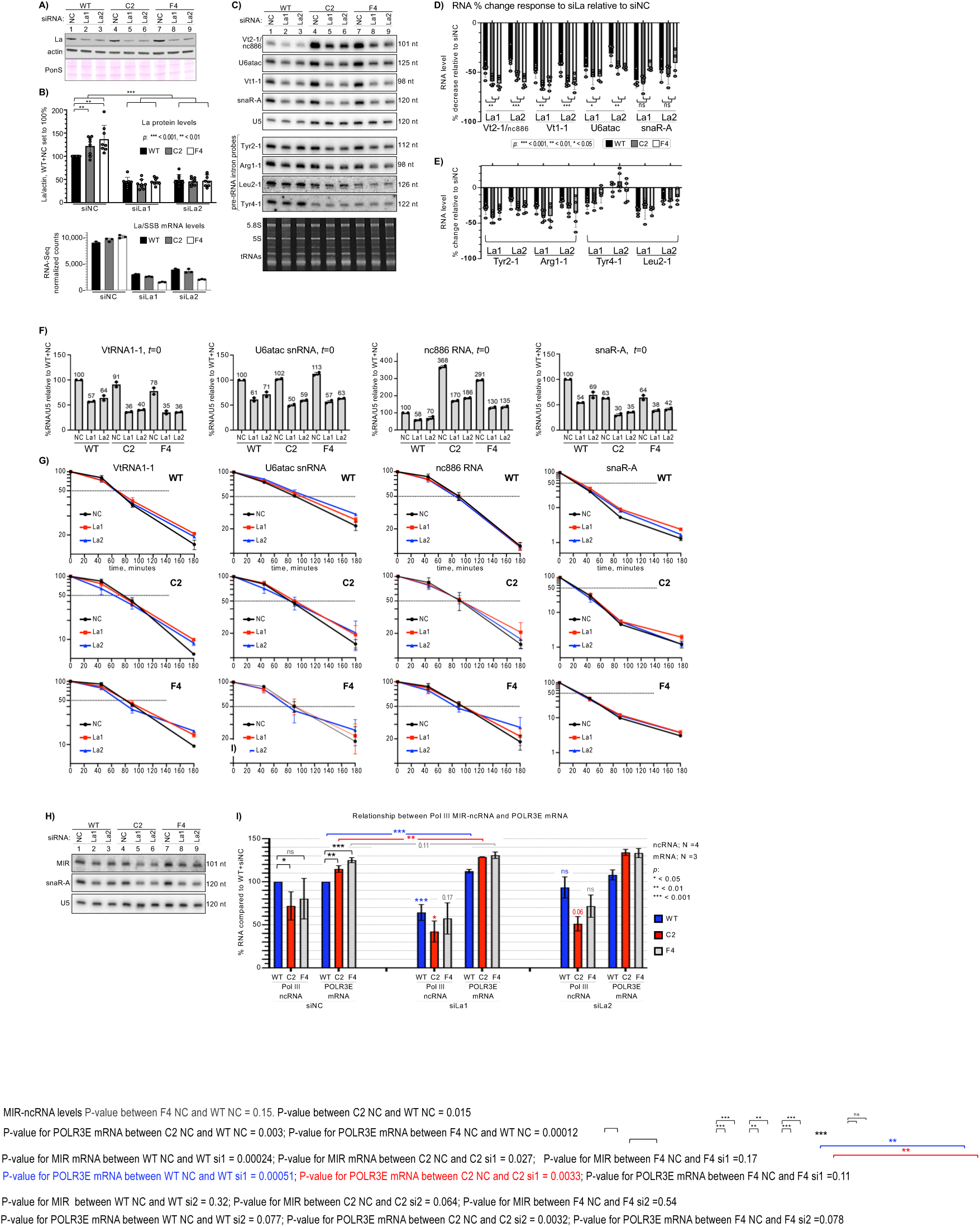
La knockdown decreases Pol III transcription products but not their half-life. **A)** Immunoblot; La1 and La2 are siRNAs directed to La/SSB mRNA; NC is negative control siRNA. Actin is a loading control. The ponceau S stained membrane prior to transfer is shown. **B) Upper panel:** Quantification of La protein levels from replicate experiments; NC in WT from each was set to 100%; error bars represent the SD. **Lower:** La/SSB mRNA levels from a triplicate RNA-Seq experiment. **C)** Northern blot of total RNAs, probed for transcripts indicated to the left. **D-E)** Quantification of repressive effects of siLa1 and siLa2 compared to siNC on ncRNA levels, in WT, C2 and F4 cells as indicated, from replicate experiments as in C; U5 used for normalization. Error bars represent the SD. *P* values were calculated using a two-tailed Student’s two-sample equal variance t-Test. **F)** Quantifications of steady state, *t* = 0, RNA levels from duplicate samples of WT, C2 and F4 cells as part of time course analyses. **G)** ActD time course decay profiles from quantification of duplicate northern data as shown for one experiment in Sup Fig S4B for the ncRNAs indicated in WT (top row), C2 (middle) and F4 (bottom) cells. **H)** Northern blot probed for *POLR3E* intron MIR-ncRNA and snaR-A ncRNA as a control for efficacy of siLa vs. siNC, indicated to the left. **I)** Bar graph showing covariation of POLR3E mRNA levels in siLa1, siLa2 and siNC (N = 3) and the POLR3E-MIR ncRNA (N = 4) as indicated. Error bars represent the SD. P values were calculated using a two-tailed Student’s two-sample equal variance t-Test. The asterisks and numbers without brackets (siLa1, siLa2) indicate p value comparisons to the color coded siNC component.

La is known to associate with a fraction of VtRNA^59^ and other ncRNAs. We asked if La-KD would alter turnover of these ncRNAs. They were analyzed at *t*=0 and after transcription inhibition **(Sup Fig S4)**. KD of the transcripts at *t*=0 was confirmed **(Fig 6F)**. Turnover profiles in siNC-cells were similar to those of untreated cells, and consistent with prior data **(Fig 6G)**^45,58,60,61^. Importantly, RNA *t*½ differences between siLa and siNC were small relative to the differences between their steady state levels at *t*=0, consistent with transient La binding^59,62,63^ whereas significant loss of stabilization was not observed. This is *in vivo* evidence that La contributes to Pol III transcription in cells, supporting previous *in vitro* data.

### A *POLR3E* auto-upregulation response to decrease in Pol III transcription increases after La-KD

Decreasing levels of Pol III transcription of a type-2 MIR-ncRNA gene that opposes the *POLR3E* promoter leads to higher levels of *POLR3E* mRNA^32^, confirmed in HEK293 cells^64,65^. *POLR3E* is regulated by Pol III in *cis* but not when the MIR-ncRNA gene is moved to other loci^32^. Thus, higher *POLR3E* mRNA in *POLR3B***^1625A>G^** cells (Fig 2) relative to WT may reflect autoregulation.

Relative to U5 loading control, siLa1-2 decreased MIR-ncRNA levels with patterns similar to snaR-A **(Fig 6H)**. In the siNC set, MIR-ncRNA levels were lower in *POLR3B***^1625A>G^**cells than in WT although more statistically significant for C2 than F4, while *POLR3E* mRNA levels were higher (**Fig 6I**). With siLa1, MIR-ncRNA decreased and *POLR3E* increased in cells relative to siNC levels **(Fig 6I)**. Again, siLa2 was less efficacious than siLa1 at decreasing expression; it minimally decreased MIR-ncRNA levels in WT cells, and *POLR3E* was also less increased **(Fig 6I)**. Nonetheless, in siLa1-treated cells, decreases in Pol III transcription upregulated *POLR3E* mRNA levels. The data support the idea that auto-upregulation of *POLR3E* mRNA contributes to the Pol III-deficiency response in *POLR3B***^1625A>G^**cells.

### La-dependent upregulation of pre-tRNA 3’-trailer-derived tRF-1s in *POLR3B*^1625A>G^ cells

tRF-1s, tRF-3s, and tRF-5s comprise major tRF classes, derived from different regions of tRNA transcripts although internal-tRFs also exist^66^. tRF-1s are 3’-trailers, formed by cleavage at the discriminator position in pre-tRNA by RNase Z/ELAC2 and Pol III termination at a U(n)U site^67^. Compared to tRF-5s and -3s, tRF-1s are produced by fewer tRNA genes^33,66^, are much more unstable due to the 5’-3’ exonuclease XRN2, yet several accumulate to much higher levels^68^. La binds 3’-U(n)U of tRF-1001 (aka tRF-U3-1) and other highly abundant tRF-1s, and promotes their accumulation^33^. tRF-1001 can promote cellular proliferation^67^.

Small-RNA sequencing (sRNAseq)^69^ was performed on C2, F4 and WT cells treated with siRNAs. Reads were sequentially mapped to custom reference genomes; miRNAs **+** other ncRNAs, mature-tRNAs, and genomic tRNAs followed by ample 3’-sequence to annex variably-positioned tRNA gene-specific terminators (Methods). Only unmapped reads from an alignment were used for the next (**Sup Fig S5A**). Replicates revealed strong correlations, and mapped fragments were of expected sizes (**Sup Fig S5B,C**). Lower relative levels of tRFs derived from mature-tRNAs in C2 and F4 cells were accompanied by higher levels of pre-tRNA-derived sRNAs as compared to WT (counts per million, CPM) **(Fig 7A**, **Sup Fig S5D)**. Mapping revealed pre-tRNA-derived sRNAs predominantly as tRF-1s, more abundant in C2 and F4 than in WT and more decreased by siLa-RNA (**Fig 7B**, **Sup Fig S5E**, **Fig 7C upper**). The mature-tRNA-derived sRNAs mapped across tRNA differently in C2 and F4 relative to WT and at lower levels (**Fig 7C lower**). Importantly, ratios of tRF-1s to mature-tRNA-tRFs were higher in *POLR3B**^1625A>G^***cells than in WT.

**Figure 7:**
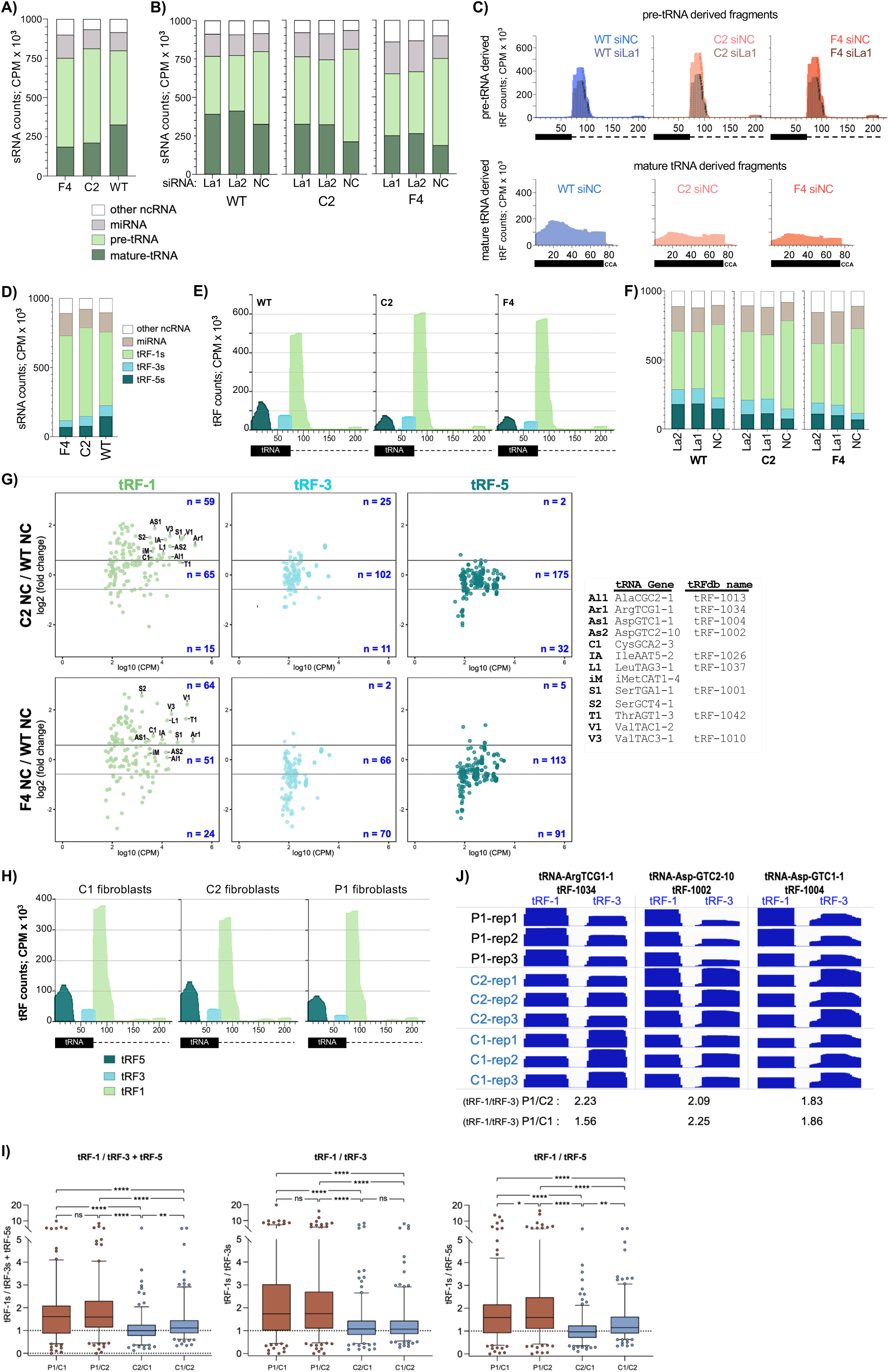
3’ trailer-derived tRF-1s differentially accumulate in *POLR3B*^1625A>G^ cells and in *POLR3B*-deficient patient cells. **A)** Distributions of four categories of small (s)RNA fragments following alignment-mapping as depicted in Sup Fig S5A, derived from mature-tRNAs, pre-tRNAs, other non-coding (nc)RNAs, and the miRNA database, in counts per million (CPM) by color code inset. See Sup Fig S5D for graph of the same data with each category separated. **B)** As in A together with data samples for La1 and La2 siRNAs directed to La/SSB mRNA and negative control siRNA, NC. See Sup Fig S5E for graph of the same data with each category separated. Note; NC samples here are the same as WT, C2 and F4 in A. **C)** Mapping of the fragments from B. Upper: the pre-tRNA-derived fragments were mapped. The black horizontal rectangle which represents the genomic sequence of the tRNA body is followed by a dotted line representing 150 nucleotides downstream. Lower: the mature-tRNA-derived fragments were mapped. The black horizontal rectangle represents mature tRNA appended with CCA. **D)** Distributions of sRNA fragments following alignment-mapping to tRF-specific reference genomes as depicted in Sup Fig S6A (see text). **E)** Mapping of the tRF-1s, tRF-3s, and tRF-5s from D. **F)** As in D along with data samples for La1 and La2 siRNAs and negative control NC siRNA. See Sup Fig S6C for graph of the same data with each category separated. Note; NC samples are the same as WT, C2 and F4 in D. **G)** MA plots of log2-fold change differences (Y-axis) in levels of miRNAs and tRFs from individual tRNA gene loci between C2 and WT (upper row) and F4 and WT (lower), and the average expression levels in log10 CPM on X-axis. tRNA source genes of individual tRF-1s validated by IGV are indicated. **H)** Mapping of tRF-5s, tRF-3s, and tRF-1s from triplicate patient (P1) and control C1 and C2 fibroblast samples following alignment-mapping using tRF-specific references (see Fig S7B). **I)** Three box and whisker plots representing triplicate normalized average CPM (Y-axis) for tRF1/tRF3 **+** tRF5, tRF-1/tRF3, and tRF-1/tRF5, left to right as indicated. Boxes encompass the medians and Q1-Q3 data quartiles, and whisker ends represent 5^th^-95^th^ percentile confidence intervals with outliers as individual datapoints. P values were calculated using 2-way ANOVA + Tukey’s multiple comparisons test. **J)** IGV representation of the intragenic paired tRF-1s and tRF-3s from the tRNA genes indicated atop each panel; tRFs-1034, tRF-1002 and tRF-1004 denote tRF-database nomenclature^66^. Three replicate samples for P1, and the two controls C1 and C2 are shown for each gene. The P1/C1 and P1/C2 values calculated from their tRF-1/tRF-3 averages of triplicate CPMs for tRF-1s and tRF-3s are shown below each tRF-1/tRF-3 pair. The IGV tracks were set to “group autoscale” with upper levels of the data ranges as follows; tRF-1034; tRF-1:128956, tRF-3:242, tRF-1002; tRF-1:15542, tRF-3:314, and tRF-1004; tRF-1:4428, tRF-3:144.

MA plots showed more miRNAs and pre-tRNA-derived fragments differentially increased than decreased in C2 and F4 relative to WT, whereas fewer mature-tRNA-derived fragments were increased and more were decreased (**Sup Fig S5F**). tRF-1001 from Ser-TGA-1-1 and tRF-1010/tRF-U3-2 from Val-TAC-3-1, reported to be dependent on La levels for accumulation^33^, were differentially increased in C2 and F4 relative to WT (**Sup Fig S5F**). Several abundant tRF-1s were more decreased by siLa in C2 and F4 cells than in WT, while a pre-tRNA-IleTAT2-3-derived non-tRF-1 fragment, whose formation is inhibited by La^70^, was differentially increased by siLa **(Sup Fig S5G**). These data support the hypothesis of an excess of La in *POLR3B***^1625A>G^** cells that arose from analysis of pre-tRNA transcription and metabolism.

The next analysis was to align and map our data using specific tRF-5s, tRF-3s and tRF-1s references (**Sup Fig S6A**). Again, replicates showed strong correlations (**Sup Fig S6B**) and confirmed that relative to WT cells, tRF-1s were higher (15-20%) in C2 and F4, again with tRF-5s more decreased than tRF-3s (**Fig 7D, E**). tRF-1s were lowered ~10% more by siLa in C2 and F4 than in WT (**Fig 7F**, **Sup Fig S6C**), consistent with downregulation of abundant tRF-1s (**Sup Fig S6D**). More tRF-1s were upregulated in C2 and F4 vs. WT, while fewer tRF-5s and tRF-3s were upregulated and more were down **(Fig 7G)**.

Small-RNAseq on P1, C1 and C2 fibroblasts yielded high reproducibility among triplicates (**Sup Fig S7A**). Similar to *POLR3B**^1625A>G^*** HEK293 cells, the *POLR3B-*variant P1 fibroblasts accumulated lower tRF-5 and tRF-3 levels while tRF-1 levels were comparable to those in C1 and C2 **(Sup Fig S7B**, **Fig 7H)**. The cumulative data and Pol III transcriptome response to *POLR3B****^1625A>G^****-*deficiency would predict that a fraction of tRNA genes would express tRF-1s that would accumulate in excess of tRF-3s at steady state. More tRF-1s accumulated than tRF-3s and tRF-5s per distinct tRNA gene in P1 than in C1 and C2 fibroblasts, for a substantial fraction of genes, with mean tRF-1/tRF-3 slightly higher than tRF-1/tRF-5 **(Fig 7I)**. MA plots showed that fibroblasts express abundant tRF-1s; some that are differentially increased in *POLR3B**^1625A>G^*** cells are differentially increased in P1 fibroblasts, albeit to different extents (**Sup Fig S7C**). We examined intragenic tRF-1/tRF-3 levels at specific tRNA genes that produce abundant tRF-1s and tRF-3s at levels substantially above background. This provided a guide for identifying an internally-controlled intragenic tRF-1/tRF-3 metric that was used to examine P1 relative to C1 and C2, which led to candidates as biomarkers. Results for three candidates, tRF-1034, tRF-1002, and tRF-1004, are shown in **Fig 7J**; the tRF-1 relative to tRF-3 levels from each tRNA gene are higher in P1 cells than in C1 and C2. The intragenic tRF-1/tRF-3 metrics for P1/C1 and P1/C2 are shown below the columns.

Our mapping was designed to capture tRF-1s that extended beyond the T1 terminator. This identified tRF-1 products of Pol III terminator readthrough, of which CysGCA2-3, iMetCAT1-4 and SerGCT4-1 were abundantly and some differentially expressed in HEK293 and P1 *POLR3-*deficiency cells (**Supp Fig S8**).

## DISCUSSION

We characterized a novel homozygous *POLR3B*:c.1625A>G;p.(Asn542Ser) variant from a patient with neurodevelopmental disease. The allele led to *POLR3B* mis-splicing and reduced POLR3B mRNA. In the absence of hypomyelination, Pol III functional deficiency in patient fibroblasts provided evidence of pathogenicity. Creating the *POLR3B**^1625A>G^*** nucleotide substitution in HEK293 cells caused the mis-splicing and associated deficiencies of *POLR3B* and Pol III activity. Although the substitution confers an Asn-542-Ser missense change in POLR3B protein, two types of experimental data indicate that this would have minimal, if any, contribution to Pol III deficiency in the cells. Consistent with this, cryo-EM structures show Asn-542 internal in Pol III with no apparent involvement in intersubunit interactions^2^. Thus, the major effect of the A>G substitution is mis-splicing with loss of *POLR3B* mRNA and protein.

The primary deficiency led to reduced levels of other POLR3 subunits and TFIIIBβ which recruits Pol III to tRNA and all type-2 ncRNA genes. We demonstrated decreased transcription output by the tRNA genes examined, to different extents. Examination of nascent pre-tRNA metabolism showed that decreases in output by Pol III in *POLR3B**^1625A>G^*** cells relative to WT was associated with their oligo(T) terminators and effects of La protein. Pol III output from all tRNA genes examined was decreased in *POLR3B**^1625A>G^***cells relative to WT, though those with 4T terminators were more affected than those with ≥5T terminators.

### POLR3B and Pol III regulation

In addition to being limiting for Pol III activity in *POLR3B**^1625A>G^*** cells, the data suggest that *POLR3B* is regulatory. Multiple POLR3 and TFIIIBβ subunits were coregulated in the cells. The largest subunits, POLR3A/RPC1 and POLR3B/RPC2, form the Pol III active center, comprised of an RNA-DNA binding site and protein elements with various roles in RNA production^2^. POLR3A protein was decreased in C2 and F4 cells, nearly as low as POLR3B (Fig 2M). Evidence of regulatory aspect of POLR3B is in its rescue of Pol III activity in *POLR3B**^1625A>G^*** cells. Though limiting, POLR3B over-expression restored type-2 ncRNAs to, but importantly not above, WT levels (Fig 3C). This suggests a model in which the cells are aptly responsive to POLR3B levels as a subunit key to Pol III regulation.

Increased *POLR3E* mRNA levels in *POLR3B**^1625A>G^***cells were accompanied by decreased levels of ncRNA from the Pol III MIR gene that opposes the *POLR3E* promoter, as expected of Pol III autoregulation^32,64,65^(Fig 6H-I). This *POLR3E* auto-upregulation system was responsive to decreasing Pol III activity upon depletion of La in *POLR3B**^1625A>G^*** and WT cells. Thus, we should consider POLR3E/RPC5 in context in these cells. While POLR3A and POLR3B proteins were much decreased and the initiation-like subunit POLR3F/RPC6 was also reduced, POLR3E encoding the termination-reinitiation subunit was maintained at WT levels, in excess of Pol III in *POLR3B**^1625A>G^***cells, by the regulatory circuit (Fig 2M).

Additionally, while active center formation is a major contribution of the second largest subunit of all cellular RNA polymerases, the Pol III-RPC2 homologs acquired a function that is specific to Pol III, in oligo(T)-mediated termination, in which POLR3E critically assists^11,12^. A region in POLR3B forms a tunnel for the nontemplate-DNA strand that restrains the terminator by oligo(T)-specific recognition. Assisted by RPC5/POLR3E bound to the RPC2/POLR3B lobe, this is required to induce pausing of elongation which is a prerequisite to RNA release^11,12^. Thus, it is plausible that POLR3B and POLR3E would be wired for regulation and related to Pol III termination-associated activity. Further, our data suggest that such wiring may enhance tRNA gene-specific transcription in a terminator-dependent and conditional manner.

We suspect that regulation in *POLR3B**^1625A>G^*** cells attempts to mitigate effects of low POLR3B levels by increasing specific activity of a fraction of Pol III, i.e., dependent on tRNA gene-specific features. Pol III reinitiation is associated with terminator length *in vitro*^51,55,71–74^, and correlated with Pol III occupancy on tRNA genes in mammalian cells^49^. Although our data suggest autoregulation of *POLR3E* is a component that supports efficient Pol III transcription of tRNA genes with ≥5T terminators, data to address this must await new experiments. Other factors reported to promote Pol III-recycling are La/SSB, PC4/SUB1, NFI proteins, and TOP1^71,72,4^.

Positive effects of La on *in vitro* Pol III transcription in some studies were not observed in others and confounded by transcript stabilization^75^ refs therein,reviewed ^4,76^. I*n vivo* data in mammalian cells are available in one publication, in which two La KO cell lines showed decreased levels of pre-tRNA and mature-tRNA^33^. A model of excess La relative to its Pol III transcript-ligands emerged from our results on preferential tRNA gene expression and pre-tRNA metabolism. Our data showed that pre-tRNAs and other short-lived Pol III transcripts decreased after La-KD with negligible effects due to transcript stabilization *in vivo*. This was corroborated using the Pol III-dependent *POLR3E*-MIR auto-regulatory system, which works by direct transcriptional interference of DNA-engaged Pol III rather than via the ncRNA product^32,64,65^. The cumulative data provide unprecedented evidence that La contributes to Pol III transcription in cells. Separately, La positively affects accumulation of certain 3’-processed Pol III products, such as tRF-1s.

### The Pol III transcriptome response

results in a new steady state profile of the ncRNAs of which La protein is a key factor. La binds U(n)U-3’-termini of nascent Pol III transcripts in a sequence-specific length-dependent manner^77^, with posttranscriptional stabilization well documented among other effects^52,57,78^, see ^5,6^. The model of excess La in *POLR3B**^1625A>G^*** cells that fit tRNA gene terminator expression was strengthened by differential accumulation of tRF-1s and an miRNA-related fragment known to be dependent on La^33,70^. As La was reported to interact with various miRNA-related components^70,79,80^, changes in its levels and its abundant ligands would expectedly impact miRNA biogenesis.

Beyond differences in tRNA gene-specific transcripts and other ncRNAs documented here, broader profiles of miRNAs and tRFs were differentially altered in *POLR3B**^1625A>G^*** cells. miRNAs and tRF-1s were increased in HEK293 *POLR3B**^1625A>G^***cells and patient fibroblasts, whereas tRF-5s and tRF-3s were lower than in control cells, consistent with lower Pol III activity. U6atac snRNA was increased in P1 fibroblasts and in *POLR3B**^1625A>G^***cells relative to controls, and although modest, small changes in U6atac can substantially affect expression of function-specific minor-intron Pol II genes^21^.

Demonstration of Pol III transcription deficiency in *POLR3* patient-derived cells would be important functional data toward genetic diagnosis, although this can be complicated. Based on our results, all intron-containing tRNA transcripts would not be markedly decreased. Thus gene-specific tRF-1 relative to tRF-3 (tRF-1/tRF-3) would best reflect a La-dependent Pol III transcriptome response to *POLR3B*-deficiency as biomarkers. Small-RNA-Seq of P1 fibroblasts confirmed over-expression of tRF-1s relative to tRF-3s compared to control cells. We illustrated how specific tRNA genes can be assessed for intragenic tRF-1/tRF-3 ratios as biomarkers for POLR3-deficiencies.

In *POLR3B**^1625A>G^*** cells, tRF-5s were disproportionately lower than tRF-3s. This was reminiscent of RNase Z/*Elac2* mutants that accumulated reduced levels of tRF-5s^81,82^. The mutants accumulated more terminator-associated, taRNAs from 3’-unprocessed pre-tRNAs reflecting a role of RNase Z/ELAC2 in La release^82,83^. How tRF-5 levels may reflect pre-tRNA 3’-end metabolism is unknown though intriguing possibilities exist. Finally, we note that the Pol III transcriptome response occurred here with La in excess of its transcripts in cells deficient in *POLR3B* whereas in cancer cells, tRF-1001 and other tRF-1s are high^66,67^ see refs in ^33^; and Pol III is usually upregulated, as is La^29,84^. Thus, it will be important to examine tRFs in cancer and other cells and to understand relationships among La, Pol III, and their substrate transcripts.

## DATA AVAILABILITY

The small RNA-seq data and other RNA-seq data generated in this study will be deposited in the GEO database.

## Abbreviations

ActD: actinomycin-D
BC200 RNA: brain cytoplasmic-200 RNA
CRISPR: clustered regularly interspaced short palindromic repeats
HEK293: human embryo kidney
HLD: hypomyelinating leukodystrophy
KD: knock-down
KO: knock-out
miRNA: micro-RNA
ncRNA: noncoding RNA
nc886: ncRNA-886/VtRNA2-1
P1: Proband
PCR: polymerase chain reaction
Pol III: RNA polymerase III
POLR3-RD: POLR3-related disorder
SNP: single nucleotide polymorphism
sRNA: small-RNA
*t*½: half-life
tRF: tRNA-fragment
U6atac: U6atac snRNA of the minor spliceosome
VtRNA: Vault-RNA

## ACKNOWLEDGMENTS

We thank the proband and his family for participating in this study, M. Davids for help with UDP variants, M. Brenowitz, I. Willis, R. Moir (Albert Einstein College of Medicine), T. Lowe (U California, Santa Cruz), M. Bayfield (York University) and Maraia Lab members for discussions. Computational analysis utilized resources of the NIH HPC Biowulf cluster (http://hpc.nih.gov).

## METHODS

### Human Subject and Whole exome sequencing

The proband in this study was enrolled in the NIH Undiagnosed Diseases Program^34,35^ consented under clinical protocol #76-HG-0238, “Diagnosis and Treatment of Patients with Inborn Errors of Metabolism or Other Genetic Disorders” approved by the Institutional Review Board of the National Human Genome Research Institute. This includes written parental informed consent to take part in the study and for photos to be used for publication, **Sup Fig S1**.

Whole exome sequencing, filtering of the identified variants and Sanger validation of top candidates were performed as described^85^. Variants were filtered based on minor allele frequency < 0.01 in publicly available databases such as ExAC and gnomAD (to include ethnicity-specific allele frequencies) and predicted to be pathogenic according to prediction tools SIFT, Polyphen, Mutation Taster, and CADD (phred score of greater than 15) (**Sup Table S3**). Variants with reported homozygotes in gnomAD or those that were in poorly aligned regions were excluded. Seven genes had homozygous variants, five of which were excluded after literature review (**Sup Materials and Methods and Sup Table S3**).

Primary dermal fibroblasts were from a forearm skin punch biopsy from the proband and cultured in Dulbecco’s Minimal Essential Media (DMEM) containing 4.5 g/L glucose supplemented with 10% heat inactivated fetal bovine serum and 1% antibiotic-antimycotic (Life Technologies, Carlsbad, CA). Control cells obtained from Coriell Institute (Camden, NJ) and ATCC (Manassas, VA) ATCC PCS-201-012).

### CRISPR/Cas9 knock-in HEK293 *POLR3B*^1625A>G^ cell lines

Cells containing homozygous *POLR3B* knock-in 1625A>G were obtained from Synthego Engineered Cells (Menlo Park) from their unedited pool of HEK293 WT cells, created to produce clones derived from independent editing events. The specific guide (sg)RNA sequence was 5’-UUUCUUGUCUUUCUUAAUGG and donor sequence 5’ GAAGAGCTCTCTTACCCAAATGTGTTTCTTGTCTTTCTTAGTGGTGGGTATATTATAGAGACATGTTGGTCCTAGATAGTCT. sgRNA was combined with *Streptococcus pyogenes* (sp)Cas9 to form ribonucleoprotein complex, mixed with the donor DNA and delivered to the cells via electroporation. Sanger sequencing was initially performed and used to calculate editing frequency. Edited cells were plated and harvested for single-cell clonal expansion. Independent clones were verified by sequencing as homozygous for c.1625G and distinguished from heterozygous clones. *POLR3B***^1625A>G^**designation for edited cells is used to distinguish that they were edited for a SNP whereas the *POLR3B* gene may differ elsewhere from the patient allele, e.g., in an intron not sequenced as part of exome sequencing.

### Cell culture and splicing analysis

For experimental investigation of potential mis-splicing in the proband, skin derived fibroblasts from the proband were obtained and maintained using standard cell culture conditions. Fibroblast RNA extraction and cDNA synthesis were performed as described^85^. Gene-specific primers were designed to flank the variant, PCR amplified, cloned into pCR4-TOPO vector (ThermoFisher), and sequenced using vector-specific primers. Cell lines obtained from Coriell (Camden, NJ) and ATCC (Manassas, VA) ATCC PCS-201-012) were used as controls.

For HEK293 cell splicing analysis, RNA was extracted from the HEK293 parent cell line and the homozygous c.1625A>G knock-in CRISPR/Cas9 clones C2 and F4 using TriPure (Sigma). cDNA was synthesized using *POLR3B* exon 18-specific primer 5’-TGTTGTGTTCGTACAGTGCAATGT and the Superscript III First-strand synthesis system (ThermoFisher). With the first strand cDNA as a template, PCR amplification was performed using the exon-14 forward primer 5’-TATCCGCACTGGGCATGATG and exon-17 reverse primer 5’-TGTGACTGCTGGCTTCTGTT. Products were visualized on agarose gel, and the bands were excised, eluted, and cloned into the pCR4-TOPO vector. After recovery from TOP10 *E. coli* transformants, the plasmid inserts were sequenced using vector-specific primers.

### Immunoblotting

After washing twice with PBS, cells were lysed in RIPA buffer (Pierce) with protease inhibitors (Sigma). Despite presence of these inhibitors, we found that most BDP1 appeared in fragmented forms, with minimal if any at predicted mass ~250 kDa. This was resolved by addition of PMSF (phenylmethylsulfonyl fluoride) to 1 mM to PBS immediately prior to use for cell washing and also immediately prior to lysis buffer, which led to appearance of ~250 kDa BDP1. Lysates were spun at 16,000 x g for 20 min at 4°C and supernatant transferred to a new tube. 20 ug total protein was taken up in 2x SDS loading buffer (Quality biological) containing 8% fresh β-mercaptoethanol. Samples were heated to 80°C for 5 min and electrophoresed on 4-12% NuPAGE Bis-Tris gels (ThermoFisher). Wet transfer was to a nitrocellulose membrane (0.45 uM pore, ThermoFisher) in a Mini-Cell (ThermoFisher) for 1 hr, 30V in Tris-Glycine buffer 10% MeOH. The membrane was blocked with PBS +5% milk for 1 hr at RT. Incubation with 1° antibody was overnight in PBS+5% milk, 0.05% NP-40 at 4°C. Li-Cor system provides use of 1° and/or 2° Abs tagged with different fluors and capture in different channels for quantification. Secondary antibody (Li-Cor) incubation was at 1:10:000 in PBS+5% milk, 0.05% NP-40 at RT for 1 hour. The membrane was washed 4x in PBS, 0.05% NP-40 for 5 min at RT and imaged on the Li-Cor Odyssey Clx infrared system; bands were quantified using Image Studio Light software (Li-Cor). Primary Abs were: anti-La (rabbit#25bleed7, raised against full length recombinant hLa) at 1:500; anti-Brf1 (Santa Cruz, sc-81405) at 1:500; anti-POLR3A (Cell Signaling, D5Y2D) at 1:1000; anti-POLR3B, rabbit (Bethyl, A301-855A-T) at 1:500; anti-POLR3F (Santa Cruz, SC-23917) at 1:250; anti-POLR3E (Bethyl, A303-708A) at 1:2000; anti-POLR2A (Sigma, 05-623, CTD4H8) at 1:2000; anti-Bdp1 (serum 2663, gift from RJ White)^86^ at 1:500; anti-TBP (Cell Signaling, #8515) at 1:1000; anti-γ-actin (Thermo Scientific, PA1-16890) at 1:5000; anti-β-actin (Abcam, ab8224) at 1:3000 (only used for Fig 2N); anti-GFP (Santa Cruz, sc-9996) at 1:1000; anti-SCRIB (Sigma, HPA023557) at 1:1000; anti-β-actin, mouse mAb (Sigma) at 1:1000 (only used for **Figure S1**).

### Cloning

*POLR3B* cDNA was made from HeLa RNA using Superscript III First-strand synthesis system (ThermoFisher) and used to amplify the ORF starting at the 2^nd^ codon. The PCR product was cloned in the BglII/BamHI sites of pFLAG-CMV2 (Sigma-Aldrich). The A1625G variant was introduced by site-directed mutagenesis using primers: 5’CCAAATGTTTCTTGTCTTTCTTAGTGGTAACATCTTAGGTGTCATTC and 5’GAATGACACCTAAGATGTTACCACTAAGAAAGACAAGAAACATTTGG. All constructs were verified by sequencing.

### Transfections

For control samples for protein analysis in fibroblasts, 5.5 × 10**^5^** HEK293 cells were seeded per well in a 6-well plate. The next day, 5 ug of empty vector or FLAG-*POLR3B* was transfected per well using lipofectamine 2000 (Invitrogen). After 24h, cells were split 1:5 into new wells. 48 h post-transfection, cells were washed in PBS and lysed with RIPA buffer (Pierce) +protease inhibitors (Roche).

For *POLR3B* rescue experiments, the HEK293 CRISPR/Cas9 clonal lines C2 and F4, and WT parent cells were transfected in 6-well plates with 1.5 ug empty vector, FLAG-*POLR3B* WT or the N542S version in combination with 100 ng eGFP plasmid as control. The next day cells were divided over multiple wells. 48h post-transfection cells were washed twice with 2 ml PBS, lysed, protein isolated using RIPA buffer (Pierce) containing protease inhibitors (Sigma) and RNA isolated with TriPure (Sigma) according to manufacturer’s instructions, however washing pellets 3 times with 75% EtOH instead of once.

### For La siRNA mediated knockdown experiments

the POLR3B***^1625A>G^*** C2 and F4, and parental WT cells were transfected in 6-well plates with a mix of 15 nM siRNA, 400 ng pUC19 carrier DNA, 100 ng eGFP plasmid (for transfection efficacy by visualization) and 5 ul Lipofectamine 2000, per well. Negative control siRNA: NC1 (IDT, 51-01-14-03), La siRNA #1: 5’-GGUCAAGUACUAAAUAUUCAGAUGA (IDT, Hs.Ri.SSB.13.1), La siRNA #2: 5’-UAGCAUUGAAUCUGCUAAGAAAUTT (IDT, Hs.Ri.SSB.13.2). After 24h, cells were split into multiple wells. 28h later (52h post transfection) the cells were washed twice with PBS followed by protein and RNA isolation as described above.

### Northern blotting

To make northern blots to study small RNAs, cultured skin fibroblasts from the proband and 2 different controls (using cells that were close in passage number to each other) from healthy males were seeded into 10 cm plates, 1.2 million cells per plate. The same procedure was followed for the HEK293 CRISPR cell lines, except that 6 million cells were seeded per well in 6-well plates. The next day, the cells were washed three times with 10 ml PBS, total RNA was isolated using 5 ml TriPure reagent (Sigma) per plate, using manufacturer’s instructions but washing the pellet 3x with 1 ml 75% EtOH instead of once. In the case of HEK cells in 6-well plates, cells were washed twice with 2 ml PBS before adding TriPure. Approximately 5 ug total RNA was separated in 10% polyacrylamide-TBE-UREA gel (ThermoFisher) using formamide loading buffer (Thermofisher, AM8547) and transferred to a positively charged nylon membrane (GeneScreen plus, PerkinElmer) using the iBlot2 dry blotting system (ThermoFisher). Similar to what has been found for other low abundance small RNAs, the U6atac and MIR from the *POLR3E* first intron were inefficiently transferred; addition of 2 ug sheared salmon sperm DNA (ThermoFisher, AM9680) in the loading buffer (Thermofisher, AM8547) greatly improved recovery including with 3 ug total RNA^87^. The membrane was UV-cross-linked then vacuum-baked at 80°C for 2 hr. It was then prehybridized in hybridization solution (6X SSC, 2X Denhardt’s, 0.5% SDS and 100 ug/ml yeast RNA) for one hour at the hybridization temperature (Ti). All probes were oligo-DNAs, listed in **Sup Table S6**. tRNA genes from which pre-tRNAs were examined are TyrGTA4-1, ArgTCT1-1, TyrGTA2-1, LeuCAA2-1, IleTAT1-1, and ArgTCT4-1. Hybridization with **^32^**P-end-labeled antisense probes was overnight at Ti. The blot was washed with 100 ml of 2X SSC, 0.1% SDS three times for 10 mins at RT. A final wash was with 20 ml at the Ti for 30 min. Blots were exposed to a phosphor screen and signals visualized using a phosphorimager (GE-Typhoon FLA 9500) and the signal intensity quantified using Multi Gauge V3.0 software (Fujifilm). Before the next hybridization, the blot was stripped in 0.1X SSC, 0.1% SDS at 80°C; the near complete removal (>95%) of **^32^**P was confirmed by phosphorimager analysis and a record of residual counts if any was noted.

For mRNA northern blots, RNA was isolated as described for RNAseq sample preparation; 5-10 ug RNA was separated in 1.8% formaldehyde agarose gel and transferred to a positively charged nylon overnight (GeneScreen plus, PerkinElmer). Subsequent steps were as for the small RNA blots.

### RNA decay

To measure RNA half-life, C2, F4 and WT cells were seeded into 6-well plates in duplicate or triplicate, so they had comparable densities the next day (between 50-80%). 5 ug/ml ActD (Sigma) was added to the media and cells were harvested at indicated times by washing quickly with 2 ml PBS per well and adding TriPure reagent (Sigma). Cells at time 0 did not receive ActD. Total RNA was purified following manufacturer’s instruction, except for washing the cell pellet 3x 1 ml 75% EtOH instead of once. For siRNA, cells were transfected, divided over multiple wells 24h later. After another 28h, the ActD timecourse was begun.

### Standardized quantification of intron-containing pre-tRNAs

The 3’-trailer probes to detect pre-tRNAs-TyrGTA4-1, ArgTCT1-1, LeuCAA2-1 and TyrGTA2-1 of comparable length were **^32^**P-5’ labeled by polynucleotide kinase and **^32^**P-γ-ATP using DNA and ATP concentrations that produce high specific activity probes. Triplicate blots were subjected to standard hybridization/incubation with probes at the incubation temperature (Ti)^88^ calculated from the formula: Tm = {16.6 log(M) + 0.41(*P*_gc_) + 81.5 − (675/*L*) − 0.65} where Ti 15°C lower than the Tm. For our applications M = 0.5, *P*gc = %G + C content, and *L* = nucleotide length of the DNA-oligo^88^. Ti was also used for standardized washing. **^32^**P-Oligo probe signal intensity/pixel/minute was recorded, corrected for by differences in probe half-life, if applicable.

### RNA purification for RNAseq

used the Maxwell 16 LEV simplyRNA purification system (Promega). C2, F4, and WT cells that were transfected with La siRNAs or negative control (NC) siRNAs in triplicate were harvested 52h post transfection (3 wells per cell type, siRNA). Cells were washed with 2 ml PBS per well. Per well, 80 ul of homogenization buffer containing thioglycerol (Maxwell 16 LEV kit) was added to lyse the cells. The lysates from 3 wells were combined and the Maxwell 16 LEV purification kit protocol was followed. The total DNase-treated RNA was eluted in 50 ul H**_2_**O per cartridge.

### RNA-seq analysis

of siLa1, siLa2 and siNC RNA treated cell lines C2, F4 and WT. RNA-Seq libraries were then constructed using TruSeq RNA Library Prep Kit v2 (Illumina, San Diego, CA) and sequenced on NovaSeq 6000 System S1 flowcell (Illumina). Approximately 40 million 100 bp paired-end reads were generated from each sample. Alignment of RNA-Seq data was performed with RNA-STAR (version 2.7.3)^89^ against the GENCODE human GRCh38 build. Gene-based read quantitation was performed using SubRead featureCounts (version 1.6.4)^90^. Read counts were normalized by Relative Log Expression (RLE) method^91^ and evaluated for differential expression using the Bioconductor package DESeq2^92^, comparing defined sample sets as biological replicates and drawing pairwise comparisons. Enrichment analysis of subsets of altered genes was performed using the Bioconductor package clusterProfiler (version 4.0)^93^

### Small-RNA-seq libraries

were prepared from 2-4 μg total RNA per sample and sequenced as described^69^. Libraries were size selected for 15–50 nt inserts and pooled for sequencing on Illumina NextSeq500. A first pass 18 nt cutoff bioinformatic filter was applied prior to further analysis.

#### Bioinformatic analysis of small-RNA-seq data

was by an ordered sequential alignment and mapping to multiple custom reference genomes, in which only unmapped reads from each alignment were used for the next. The HEK293 data were subjected to analysis by two different sets of complementary custom reference genomes, which provided quality control assurance information that neither alone could.

#### Custom genome creation and sRNA pseudoalignments

For custom reference genome set-1 (Sup Fig S5A), mature miRNAs were downloaded from miRbase (miRbase.org/download/CURRENT/mature.fa) and filtered for *Homo* sapiens (hsa) sequences. A reference genome “other non-coding RNAs” was created by downloading all sequences from GenCode (gencodegenes.org/human/gencode.v43.transcripts.fa), filtering out those with gene type “lncRNA” and “protein_coding”. For the mature-tRNA reference, the “all predictions” set of 619 human genomic tRNA sequences were downloaded from the Genomic tRNA database (GtRNAdb, *hg38-GRCh38 Dec 2013*) http://gtrnadb.ucsc.edu/genomes/eukaryota/Hsapi38/Hsapi38-displayed-gene-list.html), followed by removal of intron sequences where appropriate, and addition of 3’-terminal CCA. ‘All predictions’ includes the 429 high confidence gene set that most likely function in mRNA decoding plus others unfit in tRNA structure-function models^22^. For the pre-tRNA reference, the all predictions genomic tRNAs with introns if present and 150 nts downstream to include 3’-trailers and variably-distanced T1 terminators, were downloaded (5’-leaders were excluded). Indexing of these custom genomes were: (1) miRNAs + other ncRNAs, (2) mature tRNAs, and (3) pre-tRNAs using *bwa index* with *-a is* settings. Consecutive pseudoalignments were done using *bwa aln* using standard settings, followed by generation of .sam files using *bwa samse* using *–n 10* settings. Next, .sam files were converted into .bam files using *samtools sort.* These .bam files contain both mapped and unmapped reads. We used *samtools view* with *–uf 4* settings to obtained a .bam file with unmapped reads and converted this to a .fastq file for subsequent mapping using *samtools bam2fq*, followed by *seqtk seq –A* to write a standard .fastq file.

#### The second custom reference genome set

(Sup Fig S6A) employed the (1) miRNAs**+**other ncRNAs from above. Custom reference genomes for tRF-5s and tRF-3s were created from the first 35 nts only of the mature tRNA sequence file (tRF-5s) and only 30 nts from the 3’-ends including the appended CCAs (tRF-3s). The tRF-1 custom reference genome contained only the 150 nts downstream of the 3’-ends of the genomic tRNA sequences. The human snaR-A and VtRNA genes (including 50 nts downstream) were manually added from the NCBI nucleotide database (https://www.ncbi.nlm.nih.gov/nucleotide/) and defined as 5’ or 3’ fragments by dividing the sequences in halves. For this custom reference genome set all separate .fasta genomes were concatenated into one genome, indexed using *bwa index* with *–a is* settings. Pseudoalignments were done by *bwa aln* using standard settings, followed by generation of .sam files using *bwa samse* using *–n 10* settings. Finally, .sam files were converted into .bam files using *samtools sort*.

Raw sRNA-seq read counts were obtained from .bam files using *samtools view* with settings *–F 260* to obtain mapped read counts only. Differential expression analysis was done in R using the EdgeR package. Graphs were generated with normalized read counts as CPM with scatterplots, and read length distribution graphs generated in R (ggplot2) and bar graphs and read map position plots in GraphPad Prism. R code includes function to filter across samples. It calculates the 75^th^ percentile for raw counts for each gene/row across samples, then filters for 75^th^ percentile ≥20 CPM (consistent with tRFdb^66^). This ensures that genes with low CPM for an experimental sample and high CPM for a control sample for example are retained. MA plots used low cutoff of 50 CPM as per Wilson et al^68^. All computational analysis utilized computational resources of the NIH HPC Biowulf cluster (http://hpc.nih.gov).

### Web resources

BWA, http://bio-bwa.sourceforge.net/ dbSNP, https://www.ncbi.nlm.nih.gov/projects/SNP/ ExAC, http://exac.broadinstitute.org/ (Jan 2018) gnomAD, http://gnomad.broadinstitute.org/ SIFT, http://sift.jcvi.org/ CADD, http://cadd.gs.washington.edu/ GenBank, http://www.ncbi.nlm.nih.gov/genbank Mutation Taster, http://www.mutationtaster.org/ Polyphen-2, http://genetics.bwh.harvard.edu/pph2/ STRING, www.string-db.org OMIM, https://www.omim.org/ UCSC,hg19, http://genome.ucsc.edu/ Uniprot, http://www.uniprot.org/uniprot/Q12767 Alamut visual, http://www.interactive-biosoftware.com/alamut-visual/ tRFdb, http://genome.bioch.virginia.edu/trfdb/search.php GtRNAdb, http://gtrnadb.ucsc.edu/genomes/eukaryota/Hsapi38/Hsapi38-displayed-gene-list.html

## Supplementary Materials

### SUPPLEMENTARY FIGURE LEGENDS

**Supp Fig S1:**
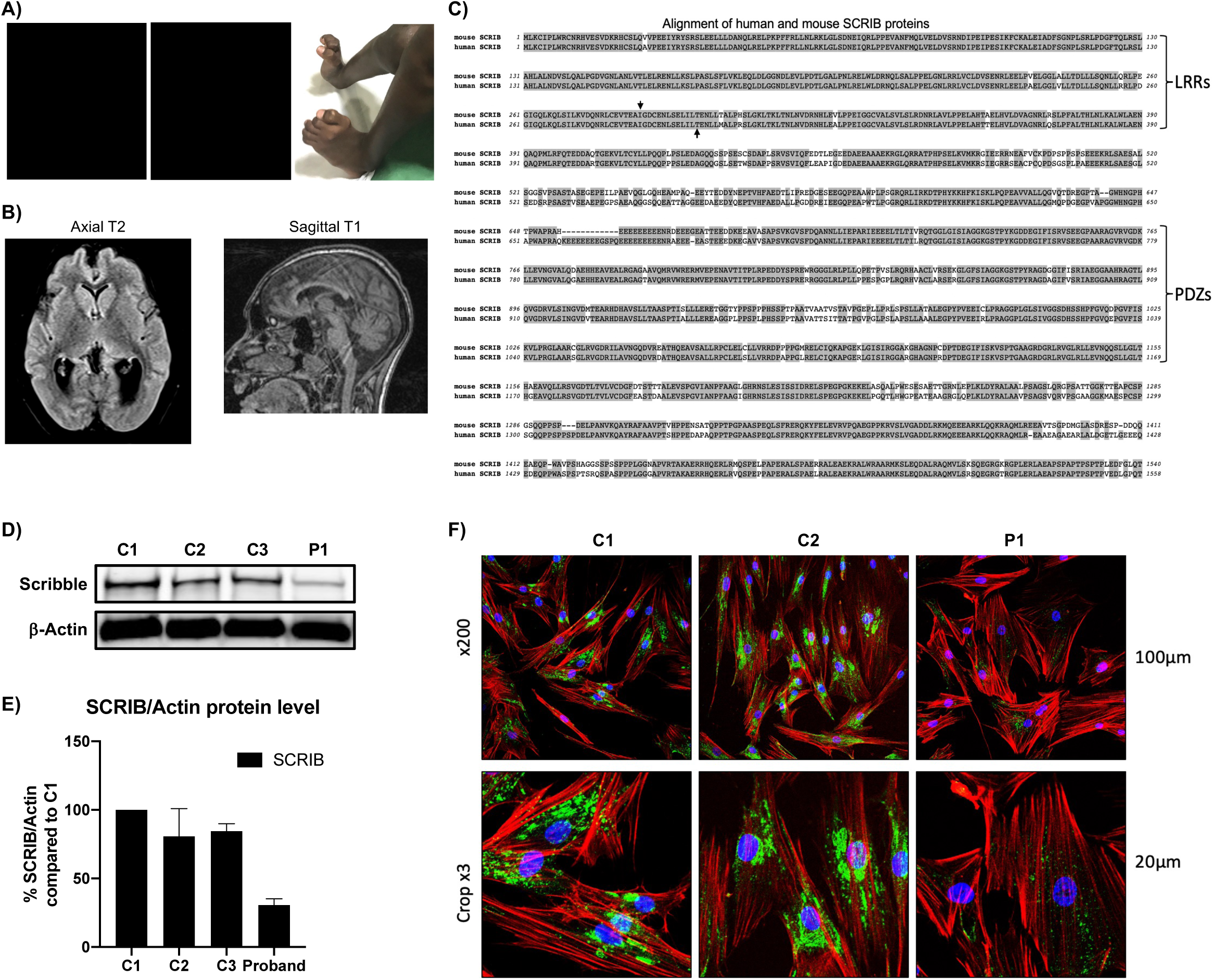
**A)** Images of the proband, P1 showing triangular facies, elevated nasal bridge, strabismus, hypodontia, hypotelorism, contractures of the toes and pes cavus. **B)** MRI images of P1 at 16 years of age showing colpocephaly and T2 hyperintensities only on the periventricular areas (axial T2 FLAIR image, left). Sagittal T1 image (right) shows frontal hypoplasia and mild thinning of corpus callosum. **C)** Sequence alignment of mouse and human SCRIB, a conserved protein with multiple leucine rich repeats (LRRs) that helps establish apico-basal cell polarity; mutations in *SCRIB* lead to neural tube defects in mice and human^1^, reviewed in ^2^. A genetic screen for developmental brain phenotypes in mice isolated multiple *SCRIB* alleles that all caused open neural tube defects in spinal cord and hindbrain; in one, the mutation was Ile285Lys^3^ (down arrow) near the *SCRIB*:c.890C>G;p.Thr297Arg in the proband, P1 (up arrow). LRRs = leucine rich repeats; PDZs = PSD-95/Disc-large/ZO-1 domains. **D-F) Examination of potential pathogenicity of the *SCRIB* variant.** The *SCRIB* mutation c.890C>G; p.Thr297Arg is associated with decreased protein levels in P1 fibroblasts. **D)** Western blot of total fibroblast protein from proband (P1) and three healthy controls (C1-3). **E)** Quantitation of western blot data; C1 was set to 100%; N = 3 technical replicates (3 blots with 3 different protein amounts loaded). Actin was used for normalization; error bars represent SD. **F)** Confocal microscopy immunofluorescence. SCRIB protein is important for the polarization of epithelial and neuronal cells, and to promote actin polymerization and cytoskeletal organization. We investigated general cell morphology here but found no evidence of disorganized actin. Actin in red, SCRIB in green. Nuclei were stained with DAPI (blue).

**Supp Figure S2:**
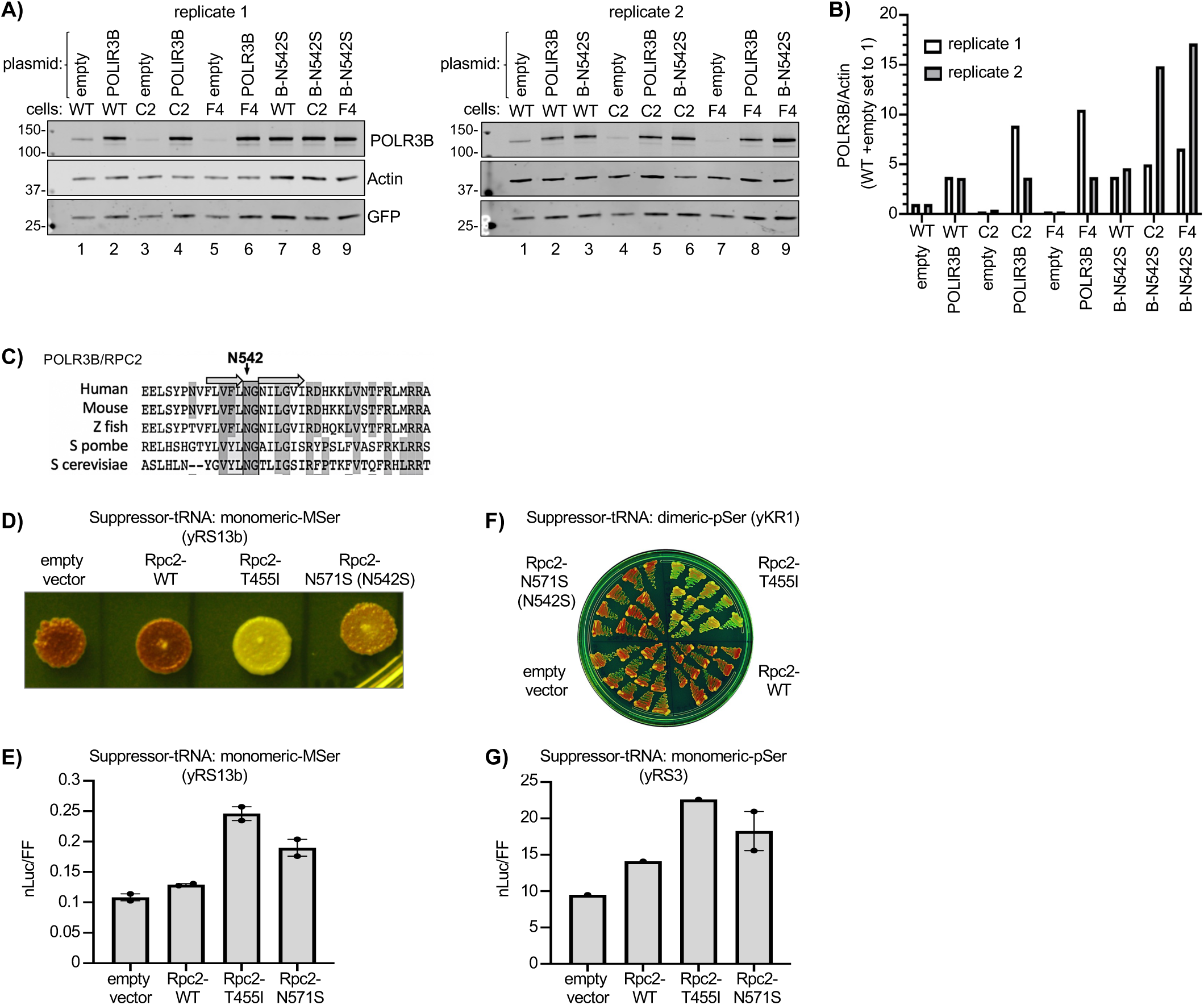
Western blots and quantification data for POLR3B ectopic expression duplicate rescue experiment and examination/assessment of potential effects of POLR3B*-N542S* on Pol III activity. **A)** Total protein isolated from cell lines WT, C2 or F4 transfected with empty vector (empty), FLAG-*POLR3B WT* (POLR3B) or FLAG-*POLR3B N542S* (B-N542S). The two western blots represent biological replicates. Antibodies used were against POLR3B, Actin and GFP. Actin was used as normalization control. **B)** Quantitation of the POLR3B signals on the western blots in A) using actin as a loading control as indicated on the Y-axis. **C-G) Examination of *S. pombe* Rpc2-*N571S* substitution corresponding to homologous conserved position in POLR3B-*N542S* for *in vivo* activity.** To determine if the *POLR3B*-SNP might alter Pol III activity independent of mis-splicing effects, we used an *S. pombe* yeast suppressor-tRNA model system^4^ for which numerous suppressor-tRNA alleles with different specific activities for suppression or designed to report on read-through past an oligo(T) terminator, are available to examine Pol III subunit and related activities reviewed in ^5^. **C)** Alignment showing that *S. pombe* Rpc2-*N571S* corresponds to conserved human POLR3B-*N542S*. **D-G)** *S. pombe* Rpc2-*N571S* was examined in three *S. pombe* strains carrying different suppressor-tRNA alleles. **D)** yRS13b carries the MSer suppressor-tRNA with modest suppression activity. In the red-white suppression (RWS) assay, spRpc2-N571S was intermediate between Rpc2-WT and Rpc2-T455I, a dual function mutant with increased transcription and terminator read-through molecular phenotypes that can be distinguished^6,7^ ^and^ ^refs^ ^therein^. **E)** Strain yRS13b also carries an integrated UGA stop codon-suppressible nanoluciferase (nLuc**^S^**) reporter and separate firefly luciferase (FFLuc) for normalization^8^. Luciferase activity generally agreed with the RWS assay in the same strain. **F)** The increased transcription and terminator read-through molecular phenotypes of the spRpc2-T455I mutant can be distinguished^6,7^ ^and^ ^refs^ ^therein^ as terminator-readthrough can be monitored specifically by the tRNA-suppressor in strain yKR1^7^. This highly sensitive RWS assay did not detect termination deficiency by Rpc2-N571S. **G)** The yRS3 strain carries a high activity pSer suppressor-tRNA, whose levels exceed the upper limit of the RWS assay^9^ which is appreciated by comparing Y-axes of panels E and G. Thus, yRS3 reports high output suppressor-tRNA gene activity (G). The cumulative data here and in HEK293 cells indicate that the missense mutation does not decrease Pol III activity. The data here suggest that when transferred to *S. pombe* Rpc2, the homologous missense substitution confers modest increase in Pol III activity.

**Supp Figure S3:**
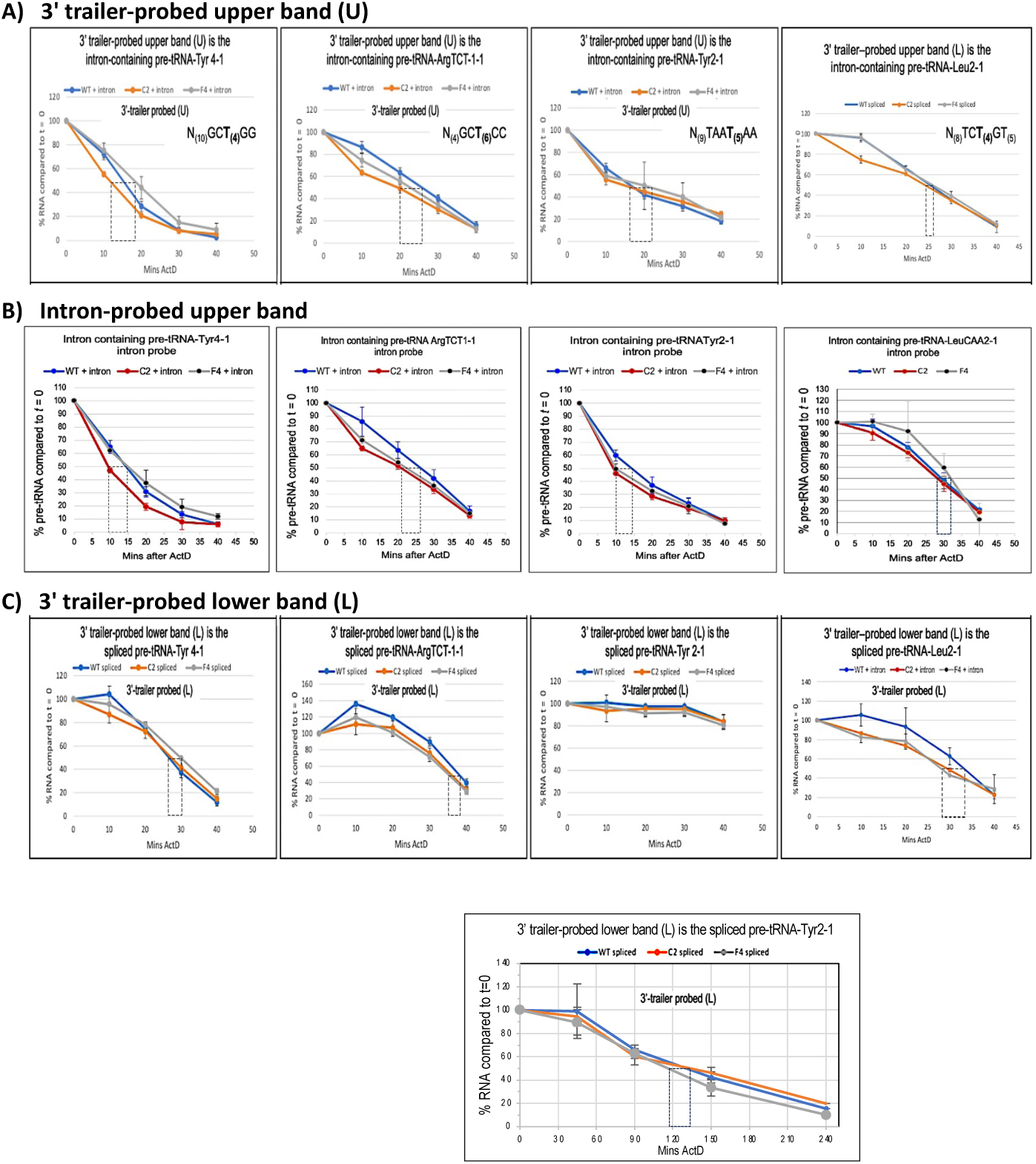
Graphic quantitative representation of RNA turnover from the gel blot data in Figure 4A. Triplicate time course northern blot data of total RNA at time zero and times after addition of actinomycin-D (ActD) as indicated on the X-axes. Cell lines WT, C2 and F4 are indicated by the colored lines; the tRNA gene names are indicated in the headers of each box. A, B and C, contain the RNA turnover profiles for the 3’-trailer probed upper bands (U) in panels i-iv of Fig 4A, the intron-probed band (panels vi-ix, Fig 4A) and the 3’-trailer probed lower bands (L, i-iv, Fig 4A). Vertical dashed-line rectangles extend from the points at which 50% of the RNAs remained (Y-axis), to the time in minutes on the X-axis, for estimating half-lives. Boxes in column A contain the 3’-trailer-terminator context sequences taken from Figure 4A.

**Supp Figure S4:**
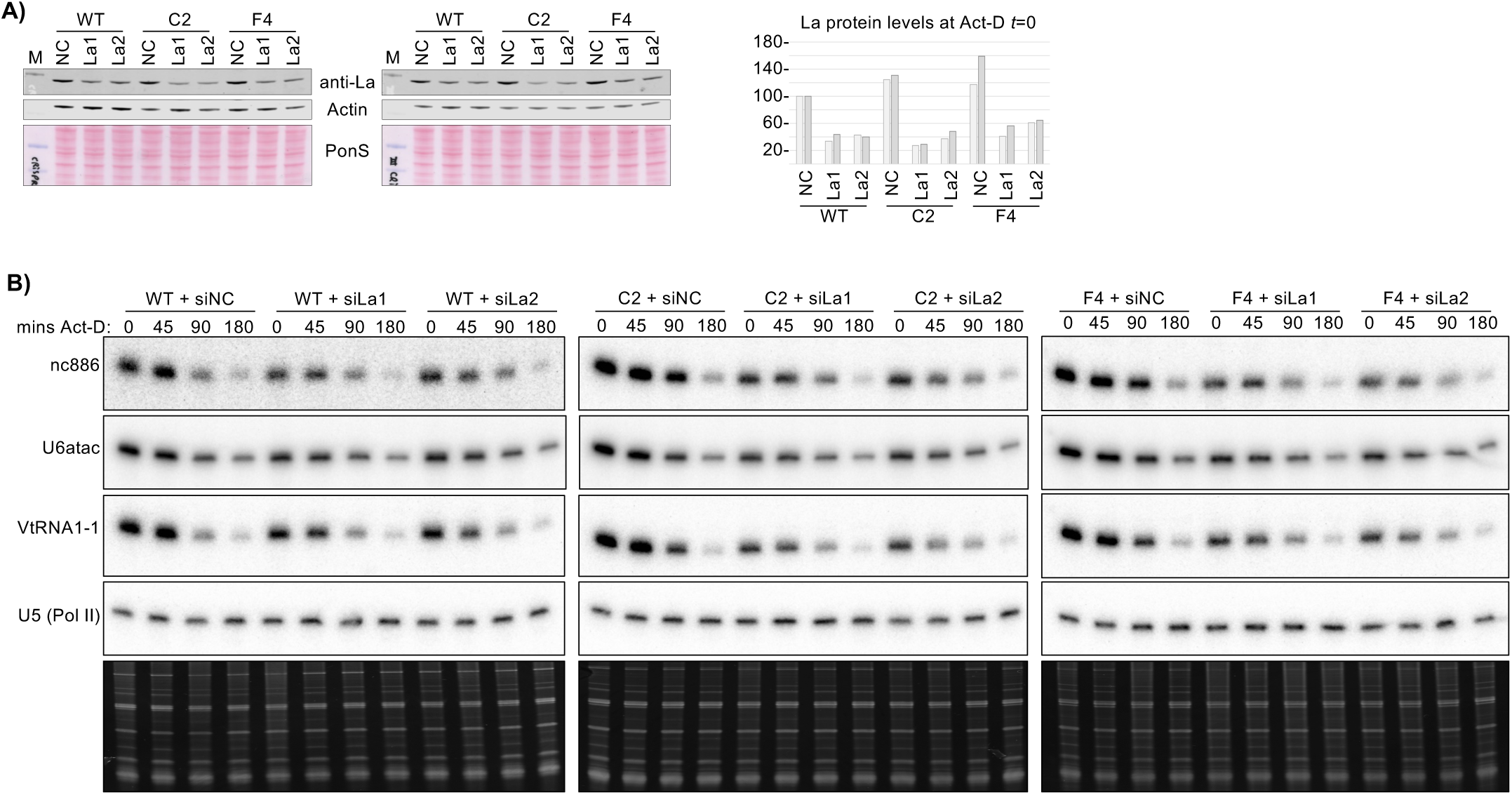
Source data corresponding to the experimental results shown in Figure 6F-G. **A)** Western blots. **B)** Quantitation of A. **C)** Northern blot RNA decay time course data of one of the biological duplicate experiments.

**Supp Figure S5:**
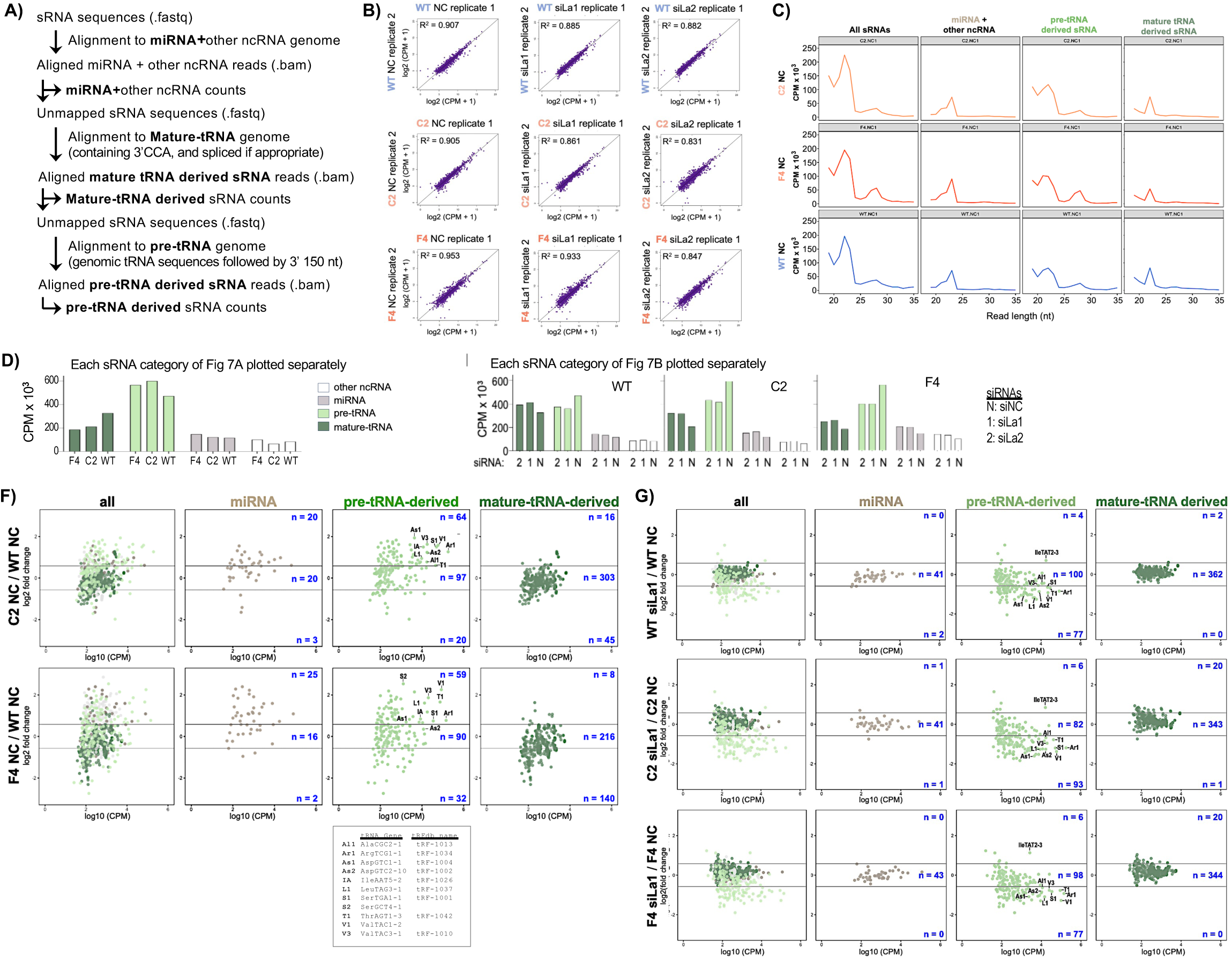
**A)** Schematic overview of the sequential alignment pipeline used for small-(s)RNA analysis. Following unique molecular identifier (UMI) deduplication and adapter trimming, sRNA sequences (in .fastq format) were sequentially aligned against three custom reference genomes comprised of i) mature miRNA sequences **+** other ncRNAs (Methods), ii) mature intron-spliced tRNAs containing 3’-CCA, and iii) genomic pre-tRNA sequences and their 3’-150 nucleotides (nt) downstream. Following each alignment only the unmapped reads were used to align against the next reference genome. **B)** Correlation plots of biological replicate samples of all mapped sRNAs from A after filtering low count reads; R^2^ values were calculated using the cor function in R (Methods). CPM: counts per million. **C)** Read length (X-axis) distribution of normalized reads (Y-axis). **D)** Amounts of normalized reads obtained for F4, C2, and WT cells, of each RNA type plotted separately, corresponding to Figure 7A, according to the color code. **E)** As for panel D but corresponding to Fig 7B. siRNAs referred to below are as follows; N: siNC, 1: siLa1, 2: siLa2. **F)** MA plots (Methods) of the data aligned and mapped following panel A and represented in Fig 7A-C show log2-fold (L2F) change differences (Y-axis) in levels of the sRNA types indicated above and average expression levels in log10 CPM on the X-axis; “all” represents a combination of sRNAs from the three reference genomes. The upper and lower rows show L2F changes between C2 and WT, and between F4 and WT, respectively. **G)** The same as F but the L2F changes are between cells treated with siLa1 or siNC as indicated for WT (upper row), C2 (middle) and F4 (lower). The tRNA source genes for some common abundant tRF-1s that have been validated by examination of integrated genome viewer (IGV) are indicated; the code legend is below.

**Supp Figure S6:**
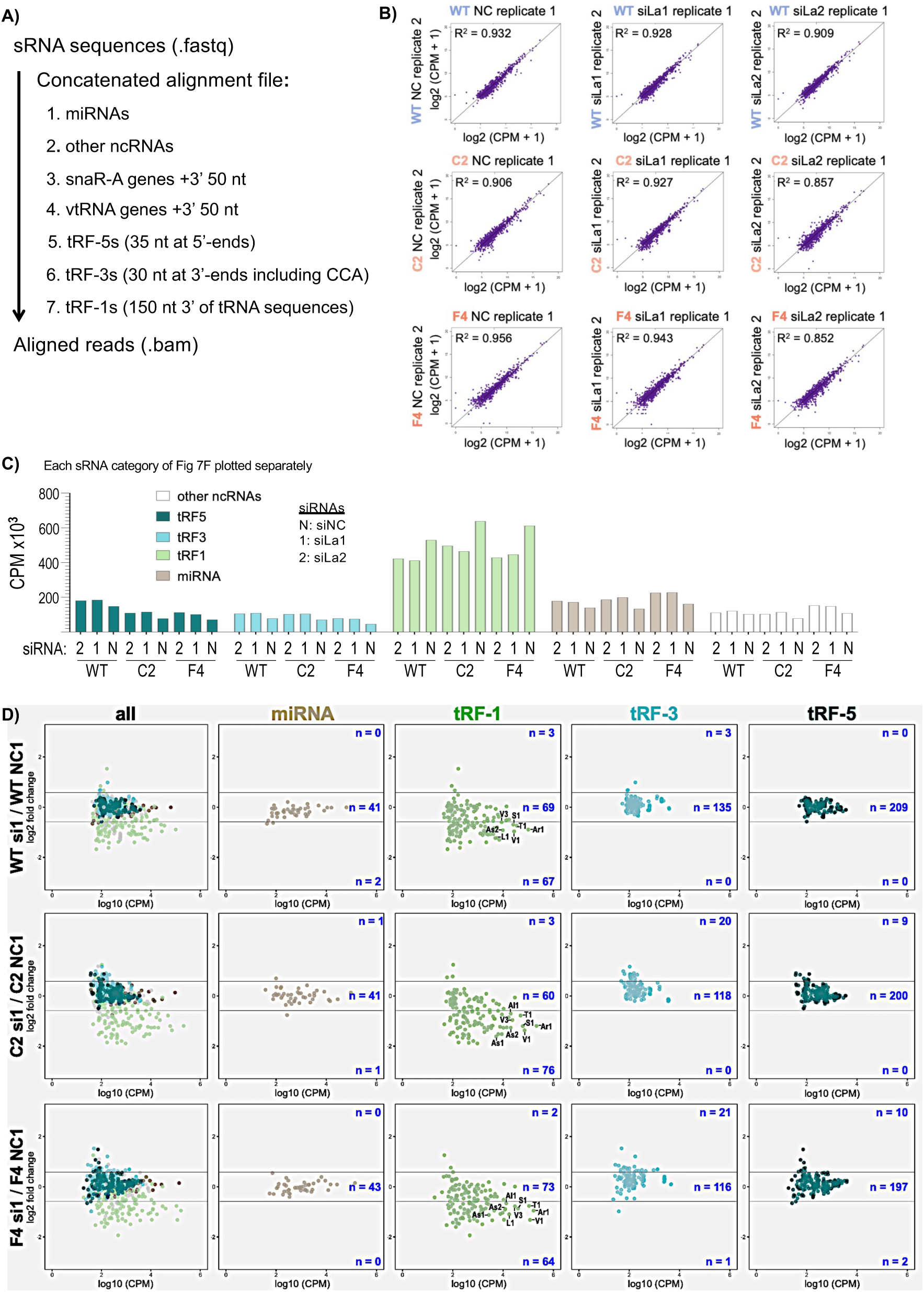
**A)** Schematic overview of alignment pipeline using custom reference genome set-2 for sRNA analysis (Methods). Following unique molecular identifier (UMI) deduplication and adaptor trimming, sRNA sequences (in .fastq format) were aligned to the custom genome file containing individual references according to the flow diagram and as described (Methods). **B)** Correlation plots of the biological replicate data after filtering low count reads from all RNA types according to figure 7F. **C)** Amounts of normalized reads obtained for F4, C2, and WT cells, of each RNA type plotted separately, corresponding to Figure 7F, according to the color code. siRNAs referred to below are as indicated. **D)** MA plots of log2-fold change differences on the Y-axis, and the average expression levels in log10 CPM on the X-axis for levels of miRNAs and the tRF-1s, tRF-3s and tRF-5s from individual tRNA genes, comparing each cell line treated with siLa1 or siNC as indicated for WT (upper row), C2 (middle) and F4 (lower row). The tRNA source genes for some common abundant tRF-1s that have been validated by by examination of integrated genome viewer (IGV) are indicated; for source gene annotations see legend under Supp Fig S5F.

**Supp Figure S7:**
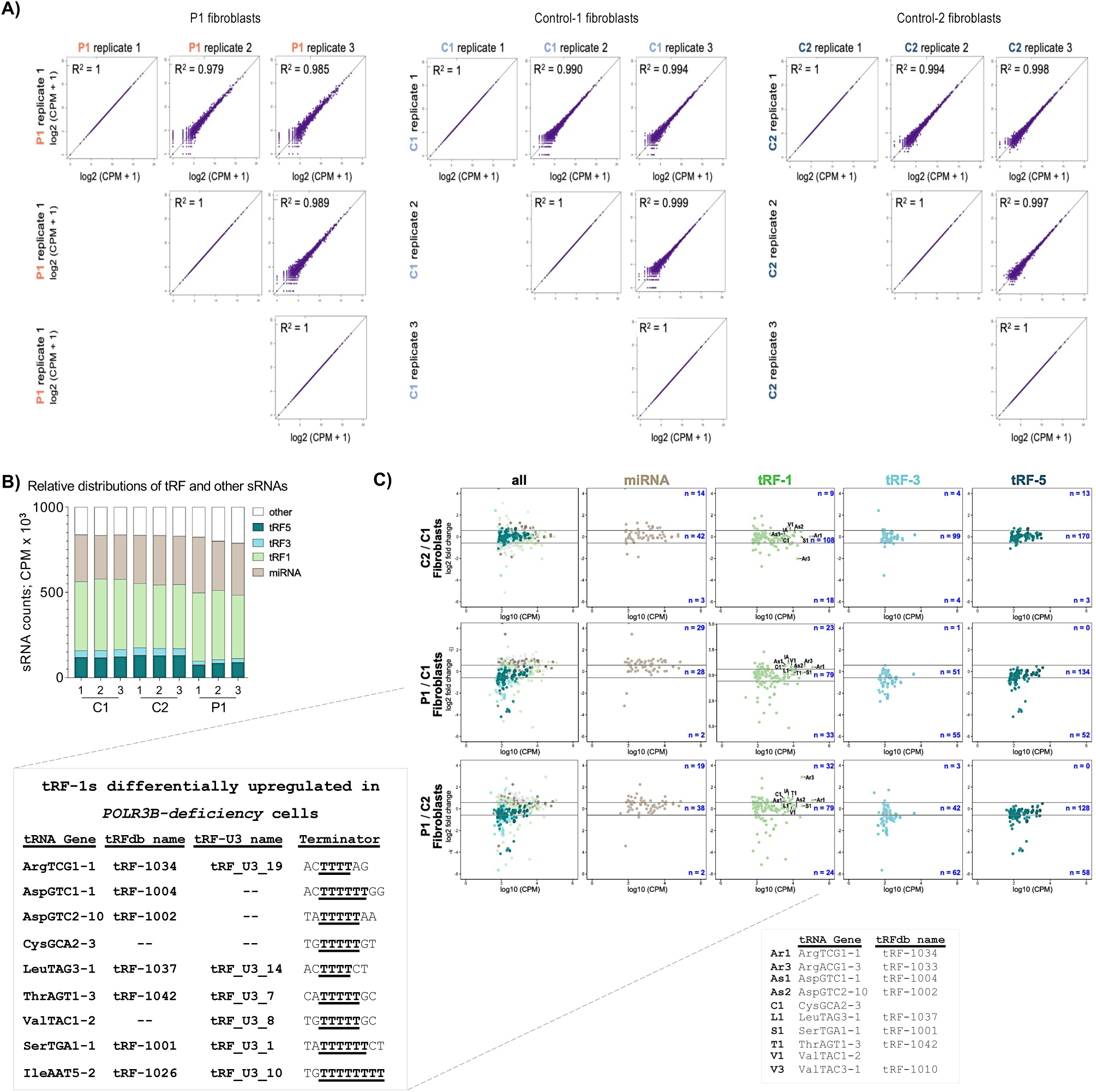
**A)** Correlation plots of biological triplicate sRNA-Seq data from patient (P1) fibroblasts, control-1 (C1) and control-2 (C2) fibroblasts after filtering low count reads from all RNA types according to Supp Fig S6A. R**^2^** was calculated using the cor function in R. **B)** Amounts of normalized reads from P1, C1 and C2 for the different RNA types according to the color-code. **C)** MA plots of log2-fold change differences (Y-axis) and the average expression levels in log10 CPM on the X-axis for levels of miRNAs and the tRF-1s, tRF-3s and tRF-5s from individual tRNA genes, comparing C2 and C1 (upper row), P1 and C1 (middle) and P1 and C2 (lower row). The tRNA source genes for some abundant tRF-1s that have been validated by integrated genome viewer (IGV) are indicated; the legend is below. An inset-like chart lists common abundant tRF-1s that are differentially upregulated in *POLR3B-deficiency* cells (see text). Their tRFdb and U3 designations if available are provided, as well as the T1 terminator motifs of the tRNA source genes.

**Supp Figure S8:**
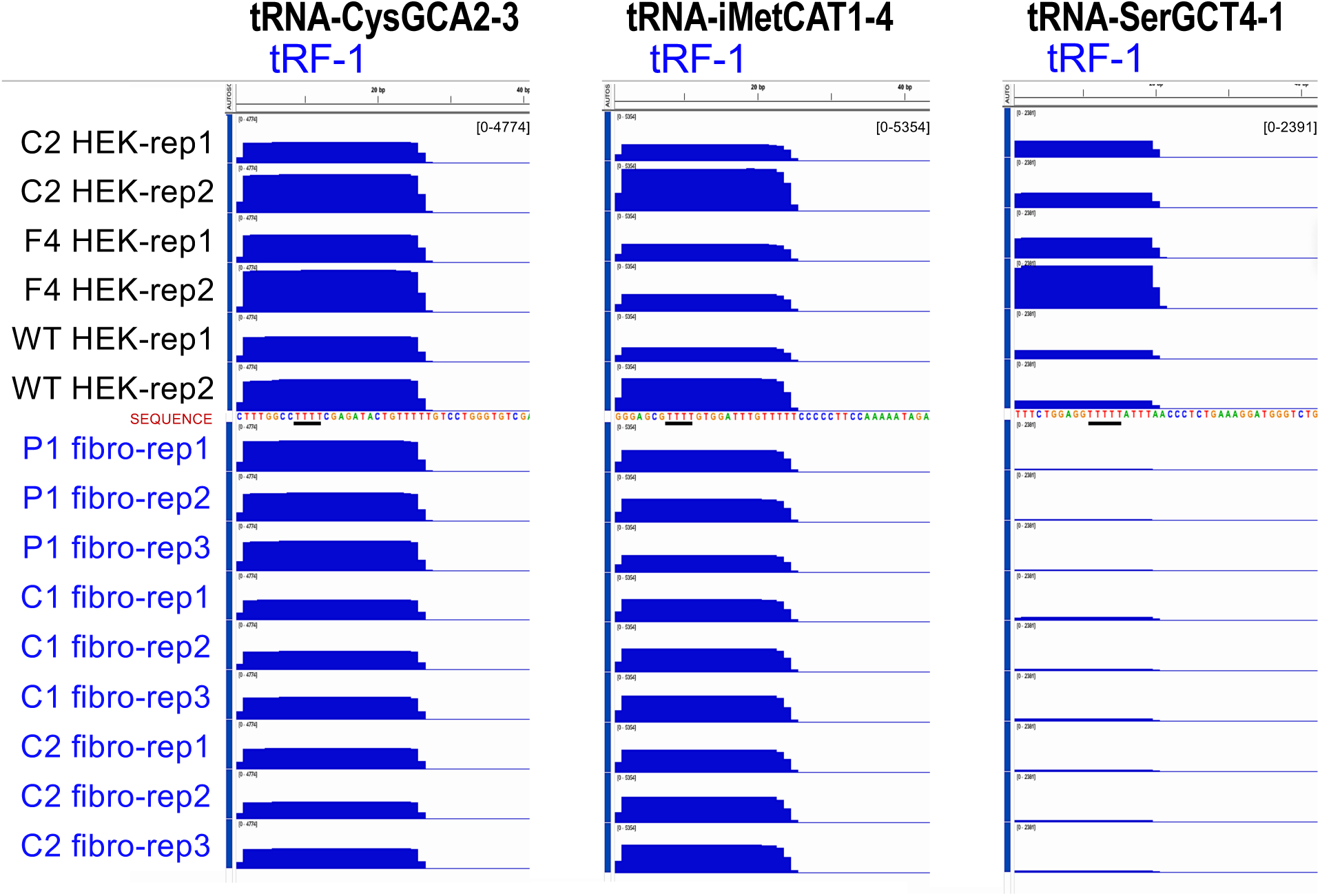
IGV representations of three high expression tRF-1s that result from Pol III terminator readthrough. Duplicate samples of HEK293 *POLR3B**^1629A>G^*** C2 and F4, and HEK293 WT below which are triplicate samples of P1, and the two controls C1 and C2 fibroblast cells. The tRNA source genes are listed above; only the sequences downstream of the genomic tRNAs are shown, with the first ≥T4 underlined. IGV tracks were set to “group autoscale” with upper levels of the CPM ranges as follows; CysGCA2-3: 4774; iMetCAT1-4: 5354, and SerGCT4-1: 2391, indicated in brackets. The HEK and fibroblast sRNA-seq libraries were processed separately by the same biochemical and bioinformatic methods.

**SUPP TABLE S1.** Features observed in the *POLR3B* variant proband (P1), male sibling (P2) and in POLR3-HLD/RD.

| Clinical feature | P1 | Sibling, P2 | POLR3-HLD (or -RD) |
| --- | --- | --- | --- |
| Short stature* | + | ND | ++ |
| Dental abnormalities* | + | ND | ++ |
| Developmental delay* | + | + | ++ |
| Thin corpus callosum* | + | ND | ++ |
| Hypomyelination* | - | - | ++ |
| Motor delay | + | + | ++ |
| Microcephaly | + | + | + |
| Colpocephaly | + | + | - |
| Tremor / ataxia | + | ND | ++ |
| Seizures | + | ND | + |
| Intellectual disability | + | ND | ++ |
| Pes cavus | + | ND | + |
| Increased deep tendon reflexes | + | ND | +<sup>1</sup> |
| Triangular facies | + | ND | +<sup>2</sup> |
| Vertical talus | + | ND | +<sup>3</sup> |
| Strabismus | + | ND | +<sup>a</sup> |
| Hypertelorism | + | ND | +<sup>b</sup> |
| Large ears | + | ND | +<sup>c</sup> |
| Fixed plantar flexion of toes | + | ND | - |
| Extension and inward rotation of foot and ankle | + | ND | - |
| Prominent eyes | + | ND | - |
| Absence of loops on thumbs and fingers | + | ND | - |

ND -not determined

**SUPP TABLE S2.** Classification of variants according to ACMG guidelines^21^.

| Gene | Variant | Strong | Moderate | Supporting | Classification |
| --- | --- | --- | --- | --- | --- |
| *SCRIB* | c.890C>G<br>(p.Thr297Arg) | - | PM2 | PP3 | Uncertain significance |
| *POLR3B* | c.1625A>G<br>(p.Asn542Ser) |  | PM2 | PP2, PP3 | ^Uncertain significance |
| *POLR3B* | c.1625A>G<br>(p.Asn542Ser) | PS3 | PM2 | PP2, PP3 | *Likely pathogenic |

^Before *in vivo* functional studies. \*After *in vivo* functionals studies were performed.PS3: Well-established *in vitro* or *in vivo* functional studies supportive of a damaging effect on the gene or gene product.
PM2: Absent from controls or at extremely low frequency if recessive in the Exome Sequencing Project, 1000 Genomes Project, or Exome Aggregation Consortium.
PP2: Missense variant in a gene with low-rate benign missense variation and which missense is common disease mechanism.
PP3: Multiple lines of computational evidence support deleterious effect on the gene/gene product (e.g., conservation, evolutionary, splicing impact).

**SUPP TABLE S3:**
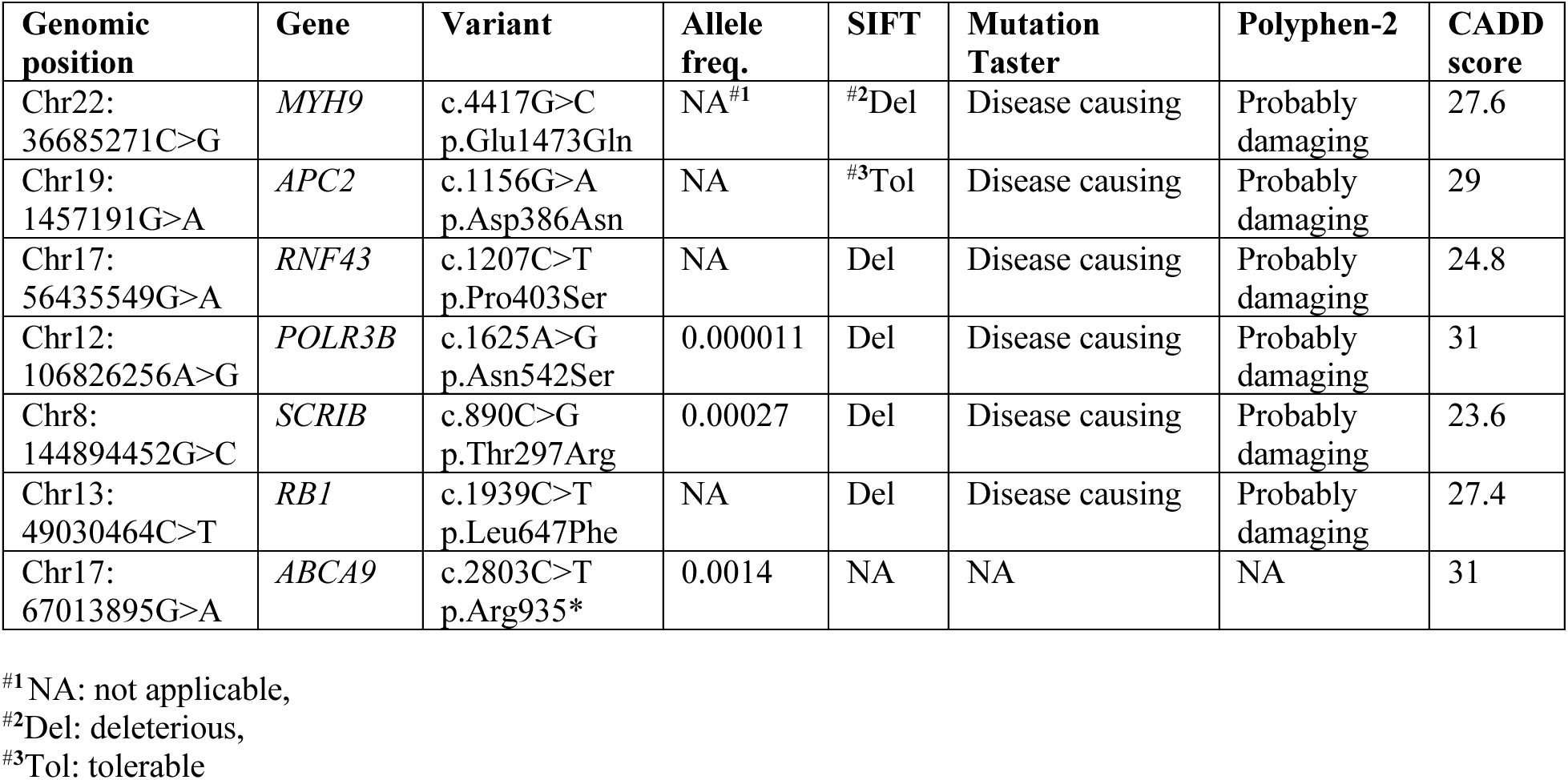
Final candidate variants identified through exome sequencing.

**SUPP TABLE S4.**
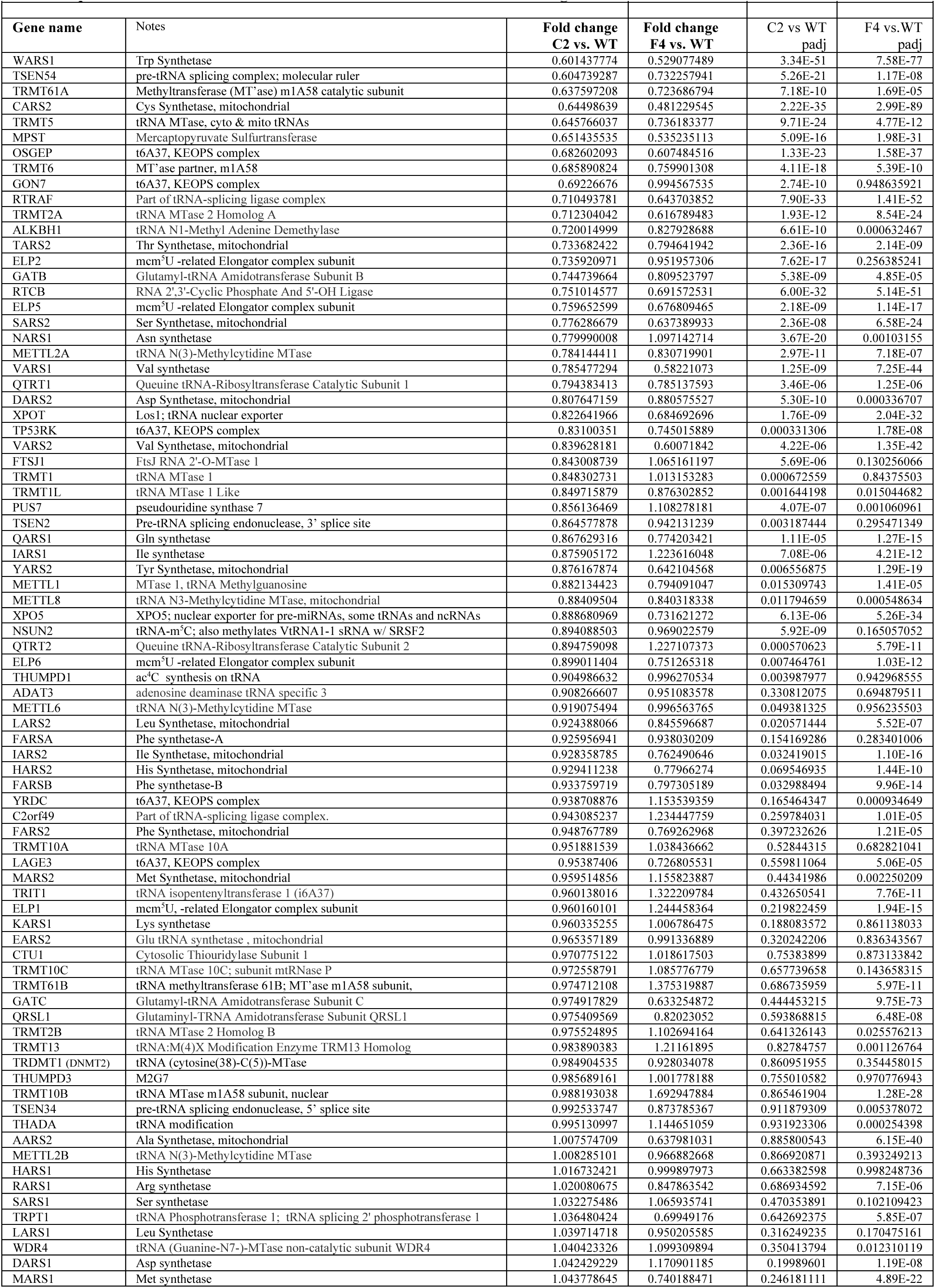
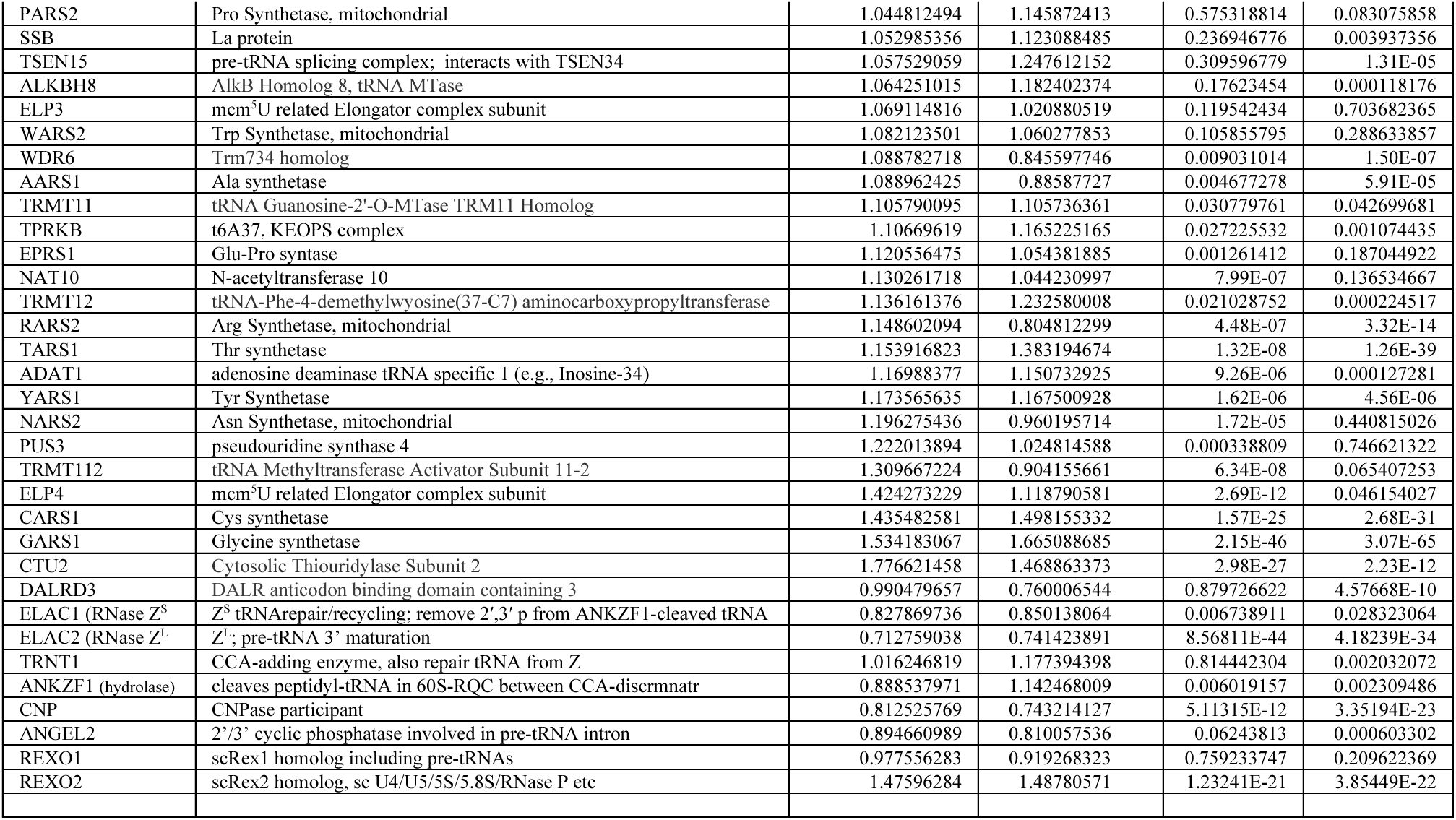
RNA-Seq normalized CPM data for tRNA metabolism and related factor genes.

**SUPP TABLE S5.**
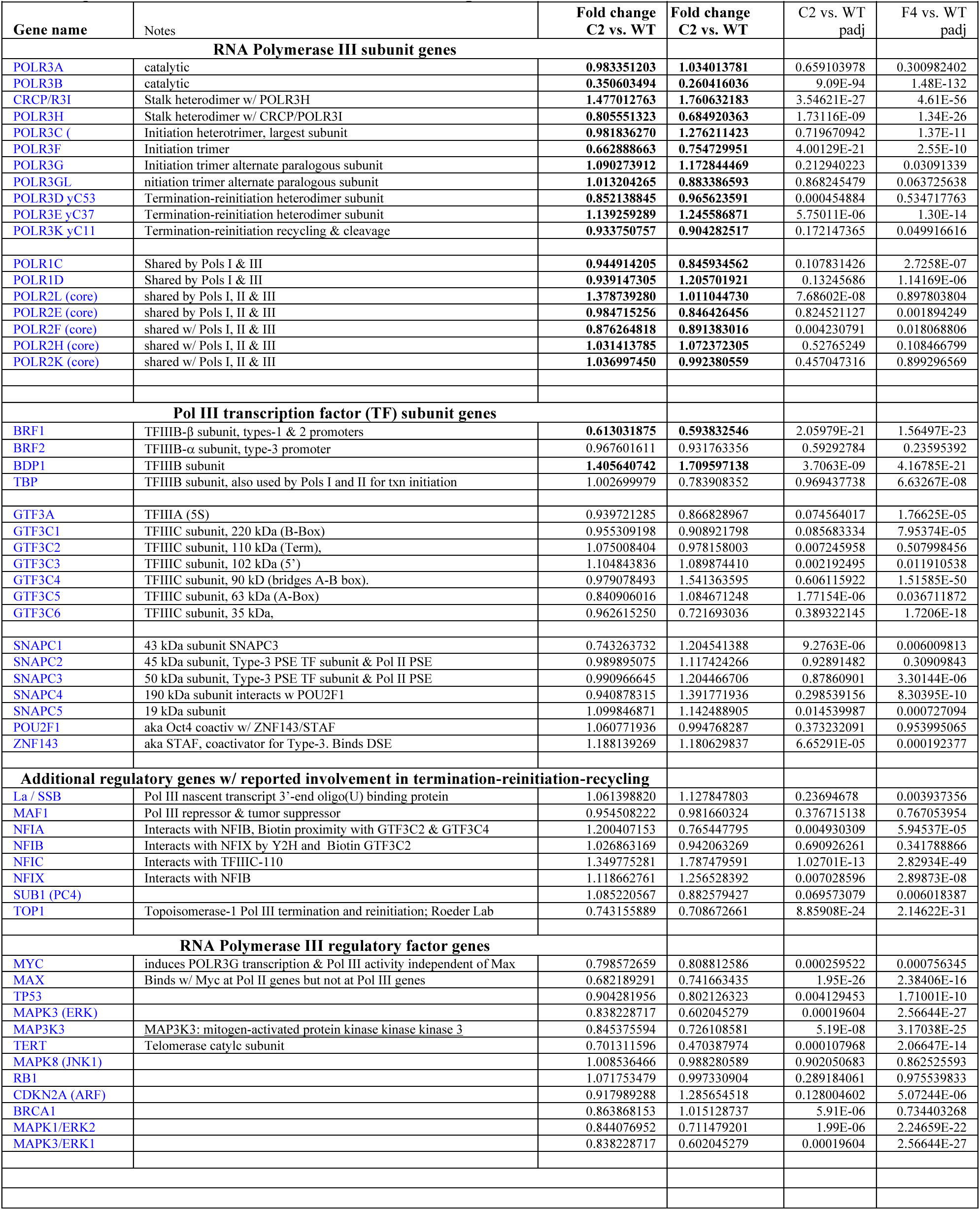
RNA-Seq normalized CPM for Pol III and related factor genes.

**SUPP TABLE S6:**
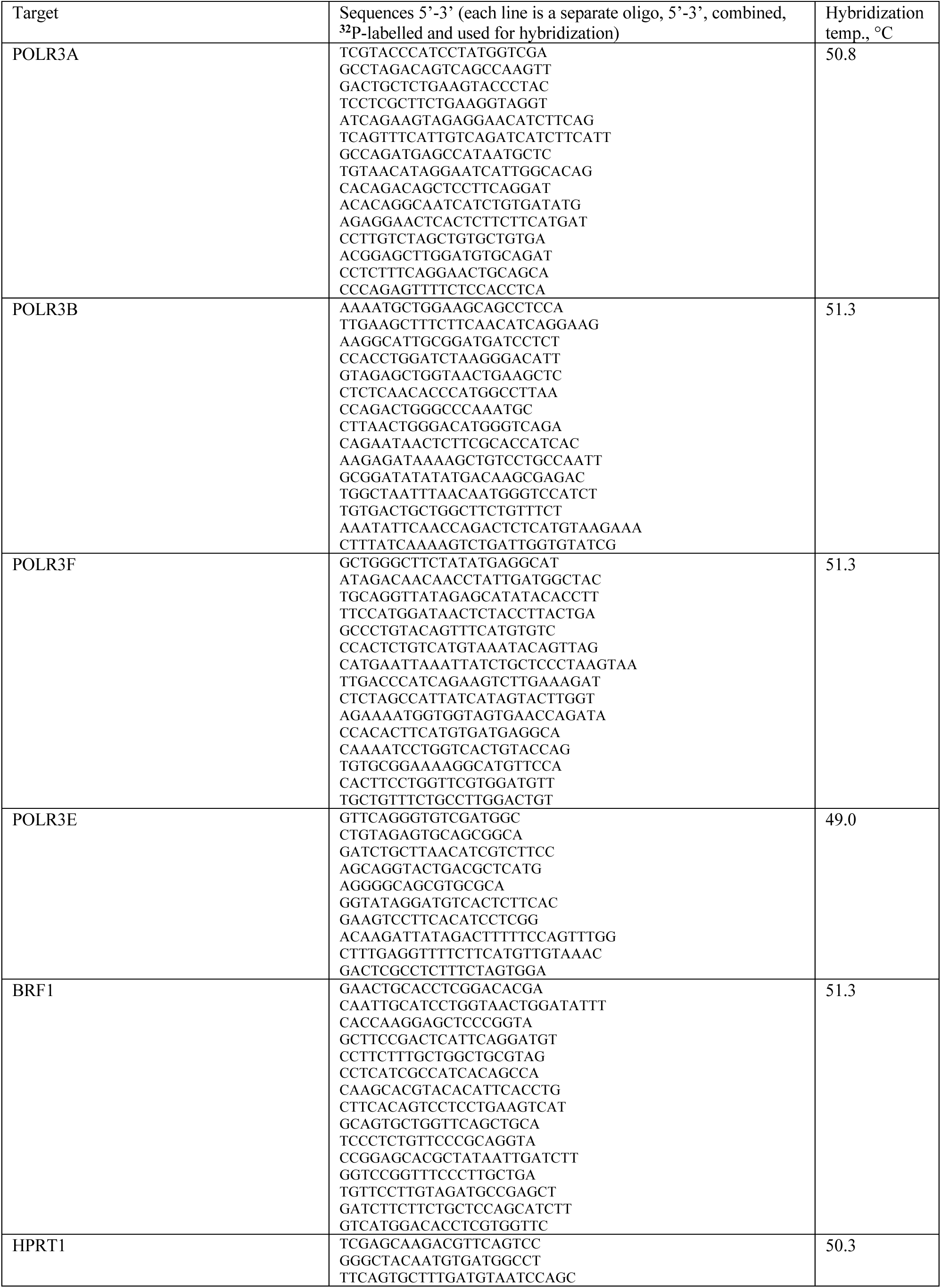
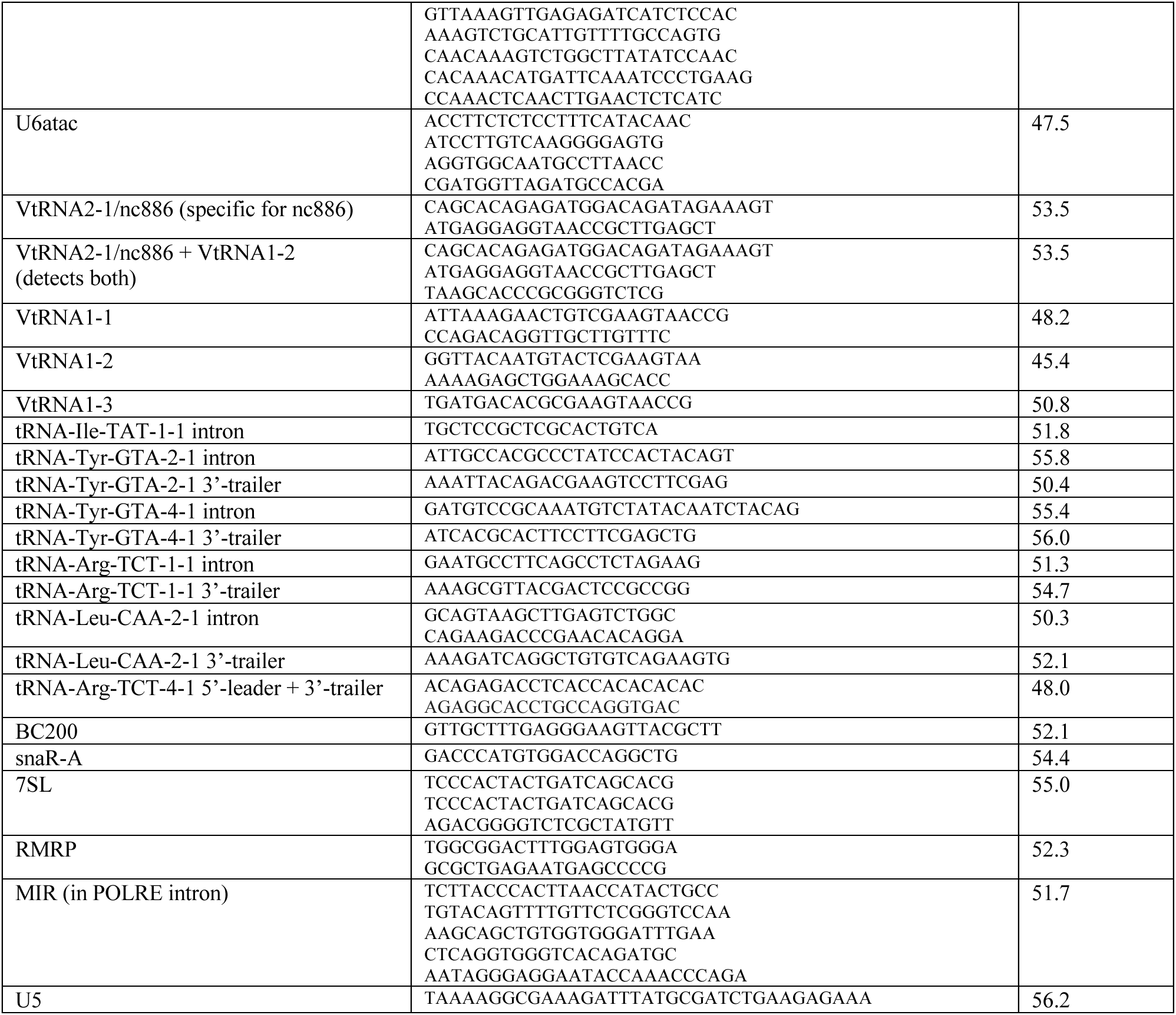
Oligo-DNA probes used for RNA detection. Multiple oligos for the same target were mixed together, equimolar, prior to end-labeling.

### SUPPLEMENTARY TEXT S1

Full clinical description of proband and younger female sibling.

The proband (P1) was a 22-year-old male (Fig 1A) from a consanguineous family, with prenatal ultrasound detection of microcephaly and a clinical history of neurological disease and developmental delay. He was a product of full-term birth, normal delivery who cried at birth. when seen in the clinic he had severe microcephaly (<1**^st^** percentile, OFC 38 cm) and motor developmental delay. He was unable to stand without support nor stand up from sitting position nor had achieved any verbal milestones. He had attention deficit hyperactive disorder, drooling of saliva and seizures from two years of age controlled by antiepileptic medication. Other features included short stature (108 cm), triangular facies, elevated nasal bridge, hypotelorism, strabismus, hypodontia, contractures of toes, pes cavus and complete absence of loops in thumbs and fingers (Fig 1B). The patient passed away in 2022 at ~24 years old.

Hearing and vision were normal at testing. CT scan showed microcephaly, a small frontal lobe with few sulci, and colpocephaly. MRI performed at age 16 years showed bilateral frontal hypoplasia, colpocephaly, moderate thinning of the corpus callosum, and completed myelination (Fig 1C).

The younger male sibling (P2) expired at age 10 with a history of a more severe phenotype. Antenatal USG showed microcephaly. CT showed microcephaly, small frontal lobe with few sulci, bifrontal subdural hygroma, colpocephaly, few calcifications around the left temporal lope and dilated lateral ventricles.

### SUPPLEMENTARY MATERIALS AND METHODS

#### Final sequence candidate variant filtering

Five of the seven genes found with homozygous variants by exome analysis were excluded after review of available literature as summarized here. *ABCA9* encodes an ATP-binding cassette (ABC) transporter involved with translocation of various substrates across membranes that is poorly expressed in nervous system tissues. *ABCA9* variants are not associated with human disease; deletion of this gene in mice leads to reduced anxiety. *RB1* (retinoblastoma transcriptional corepressor-1) is a tumor suppressor gene. Somatic mutations in *RB1* are associated with various specific types of cancer (MIMs 180200, 109800, 259500 and <u>182280</u>, none of which were detected in the proband. *MYH9* codes for non-muscle myosin heavy chain that contributes to cytoskeleton reorganization. Individuals with variants in *MYH9* have autosomal dominant deafness (MIM 603622) that can be associated with macrothrombocytopenia and granulocyte inclusions (MIM 155100). Our proband does not present with deafness. Variants in *APC2* are associated with Sotos syndrome (MIM 617169) and complex cortical dysplasia (MIM 618677), neither of which is a phenotypic match with our proband. Nonsense heterozygous germline variants in *RNF43* have been implicated in polyposis cancer syndrome (MIM 617108), while truncating or inactivating variants were identified in colorectal adenocarcinomas. No neurologic symptoms were detected in adults with *RNF43* variants.

#### Immunofluorescence/immunostaining of SCRIB protein was as described^22^

using primary anti-SCRIB, rabbit polyclonal (Sigma, HPA023557) at 1:200 and Alexafluor 488 conjugated Goat anti-Rabbit (ThermoFisher) as secondary antibody. Phalloidin was used for actin staining. Images were visualized and captured on a Zeiss LSM700 confocal laser-scanning microscope using a 20X objective and analyzed using Zen black 2012 LSM software (Carl Zeiss Microscopy GmbH, Jena, Germany).

#### tRNA mediated suppression (TMS) in *S. pombe*

The pRep4X plasmid-mediated expression in *S. pombe* harboring *ade6-704* and integrated suppressor-tRNA genes was as described^5^. *S. pombe* Rpc2-WT and Rpc2-T455I were published^6,7^; Rpc2-N571S was made by site-directed mutagenesis using Q5 Site-Directed Mutagenesis (NEB) and primers: 5’-GGTTTATTTATCTGGTGCTATTTTAGGTATTAGC and 5’AAGTAGGTACCATGGCTG. All constructs were verified by sequencing.

The Mser and pSer suppressor tRNA alleles exhibit intrinsically different specific activities for suppression, independent of terminator length. Strain yRS13b (h-*ade6-704* ura4-D18 *leu1*-32:tRNAmSer5T-*leu1**^+^***nLuc**^S1^**Kan**^R^**FLuc) was transformed with pRep4X containing Rpc2, derivatives or empty vector and plated on minimal media lacking uracil with adenine at 10 mg/L. Representative transformant colonies were spotted onto the same plates and grown at 32°C for color development. The same transformant colonies used for the above red-white spotting assay were grown overnight in liquid media lacking uracil, diluted then grown to OD600 of 0.5 in 25 ml media. Cells were harvested, and lysates were made into Glo buffer (Promega). Nano-Glo Dual-Luciferase assays were carried out according to the instructions (Promega). The same was done for yRS6 (h-*ade6-704* ura4-D18 *leu1*-32:tRNApSer7T-*leu1**^+^***nLuc**^S1^**Kan**^R^**FLuc). The nLuc**^S1^**Kan**^R^**FLuc inserted on *S. pombe* chromosome I was described^8^; the suppressor tRNA gene is at the *leu1* locus^5^. The nLuc**^S1^** indicates it is opal-suppressible, with ser-TGA codon-39 recognized by suppressor-tRNA *sup3-e* and derivatives^5^. The yKR1 strain was described^7^.

## REFERENCES

1. Kessler, A.C. and Maraia, R.J. (2021) The nuclear and cytoplasmic activities of RNA polymerase III, and an evolving transcriptome for surveillance. Nucleic Acids Res, 49, 12017–12034.

2. Girbig, M., Misiaszek, A.D., Vorländer, M.K., Lafita, A., Grötsch, H., Baudin, F., Bateman, A. and Müller, C.W. (2021) Cryo-EM structures of human RNA polymerase III in its unbound and transcribing states. Nat Struct Mol Biol, 28, 210–219.

3. Willis, I.M. and Moir, R.D. (2018) Signaling to and from the RNA Polymerase III Transcription and Processing Machinery. Annu Rev Biochem, 87, 75–100.

4. Schramm, L. and Hernandez, N. (2002) Recruitment of RNA polymerase III to its target promoters. Genes Dev, 16, 2593–2620.

5. Blewett, N.H. and Maraia, R.J. (2018) La involvement in tRNA and other RNA processing events including differences among yeast and other eukaryotes. Biochim Biophys Acta, 1861, 361–372.

6. Maraia, R.J., Mattijssen, S., Cruz-Gallardo, I. and Conte, M.R. (2017) The LARPs, La and related RNA-binding proteins: Structures, functions and evolving perspectives. WIREs RNA, e1430. doi: 10.1002/wrna.1430.

7. Maraia, R.J. and Lamichhane, T.N. (2011) 3’ processing of eukaryotic precursor tRNAs. WIRES RNA, 2, 362–375.

8. Hamada, M., Sakulich, A.L., Koduru, S.B. and Maraia, R. (2000) Transcription termination by RNA polymerase III in fission yeast: A genetic and biochemically tractable model system. J Biol Chem, 275, 29076–29081.

9. Braglia, P., Percudani, R. and Dieci, G. (2005) Sequence context effects on oligo(dT) termination signal recognition by Saccharomyces cerevisiae RNA polymerase III. J Biol Chem, 280, 19551–19562.

10. Huang, Y. and Maraia, R.J. (2001) Comparison of the RNA polymerase III transcription machinery in S. pombe, S. cerevisiae and humans (Review). Nucl. Acids Res, 29, 2675–2690.

11. Hou, H., Li, Y., Wang, M., Liu, A., Yu, Z., Chen, K., Zhao, D. and Xu, Y. (2021) Structural insights into RNA polymerase III-mediated transcription termination through trapping poly-deoxythymidine. Nat Commun, 12, 6135.

12. Girbig, M., Xie, J., Grötsch, H., Libri, D., Porrua, O. and Müller, C.W. (2022) Architecture of the yeast Pol III pre-termination complex and pausing mechanism on poly-dT termination signals. Cell Report, Sep 6;40(10):111316.

13. Lata, E., Choquet, K., Sagliocco, F., Brais, B., Bernard, G. and Teichmann, M. (2021) RNA Polymerase III Subunit Mutations in Genetic Diseases. Front Mol Biosci, 8, 696438.

14. Wilson, B. and Dutta, A. (2022) Function and Therapeutic Implications of tRNA Derived Small RNAs. Front Mol Biosci, 9, 888424.

15. Persson, H., Kvist, A., Vallon-Christersson, J., Medstrand, P., Borg, A. and Rovira, C. (2009) The non-coding RNA of the multidrug resistance-linked vault particle encodes multiple regulatory small RNAs. Nat Cell Biol, 11, 1268–1271.

16. Ahn, J.H., Lee, H.S., Lee, J.S., Lee, Y.S., Park, J.L., Kim, S.Y., Hwang, J.A., Kunkeaw, N., Jung, S.Y., Kim, T.J. et al. (2018) nc886 is induced by TGF-beta and suppresses the microRNA pathway in ovarian cancer. Nat Commun, 9, 1166.

17. Sajini, A.A., Choudhury, N.R., Wagner, R.E., Bornelöv, S., Selmi, T., Spanos, C., Dietmann, S., Rappsilber, J., Michlewski, G. and Frye, M. (2019) Loss of 5-methylcytosine alters the biogenesis of vault-derived small RNAs to coordinate epidermal differentiation. Nat Commun, 10, 2550.

18. Golec, E., Lind, L., Qayyum, M., Blom, A.M. and King, B.C. (2019) The Noncoding RNA nc886 Regulates PKR Signaling and Cytokine Production in Human Cells. J Immunol, 202, 131–141.

19. Fort, R.S. and Duhagon, M.A. (2021) Pan-cancer chromatin analysis of the human vtRNA genes uncovers their association with cancer biology. F1000Res, 10, 182.

20. Stribling, D., Lei, Y., Guardia, C.M., Li, L., Fields, C.J., Nowialis, P., Opavsky, R., Renne, R. and Xie, M. (2021) A noncanonical microRNA derived from the snaR-A noncoding RNA targets a metastasis inhibitor. RNA, 27, 694–709.

21. El Marabti, E., Malek, J. and Younis, I. (2021) Minor Intron Splicing from Basic Science to Disease. Int J Mol Sci, 22.

22. Chan, P.P., Lin, B.Y., Mak, A.J. and Lowe, T.M. (2021) tRNAscan-SE 2.0: improved detection and functional classification of transfer RNA genes. Nucleic Acids Res, 49, 9077–9096.

23. Van Bortle, K., Marciano, D.P., Liu, Q., Chou, T., Lipchik, A.M., Gollapudi, S., Geller, B.S., Monte, E., Kamakaka, R.T. and Snyder, M.P. (2022) A cancer-associated RNA polymerase III identity drives robust transcription and expression of snaR-A noncoding RNA. Nat Commun, 13, 3007.

24. Bernard, G., Chouery, E., Putorti, M.L., Tetreault, M., Takanohashi, A., Carosso, G., Clement, I., Boespflug-Tanguy, O., Rodriguez, D., Delague, V. et al. (2011) Mutations of POLR3A encoding a catalytic subunit of RNA polymerase Pol III cause a recessive hypomyelinating leukodystrophy. Am J Hum Genet, 89, 415–423.

25. Tetreault, M., Choquet, K., Orcesi, S., Tonduti, D., Balottin, U., Teichmann, M., Fribourg, S., Schiffmann, R., Brais, B., Vanderver, A. et al. (2011) Recessive mutations in POLR3B, encoding the second largest subunit of Pol III, cause a rare hypomyelinating leukodystrophy. Am J Hum Genet, 89, 652–655.

26. Saitsu, H., Osaka, H., Sasaki, M., Takanashi, J., Hamada, K., Yamashita, A., Shibayama, H., Shiina, M., Kondo, Y., Nishiyama, K. et al. (2011) Mutations in POLR3A and POLR3B encoding RNA Polymerase III subunits cause an autosomal-recessive hypomyelinating leukoencephalopathy. Am J Hum Genet, 89, 644–651.

27. Bernard, G. and Vanderver, A. (2017) In Pagon, R. A., Bird, T. D., Dolan, C. R., Stephens, K. and Adam, M. P. (eds.), GeneReviews® 1993-2021. University of Washington, Seattle, Vol. Last Update: 2017.

28. Perrier, S., Gauquelin, L., Fallet-Bianco, C., Dishop, M.K., Michell-Robinson, M.A., Tran, L.T., Guerrero, K., Darbelli, L., Srour, M., Petrecca, K. et al. (2020) Expanding the phenotypic and molecular spectrum of RNA polymerase III-related leukodystrophy. Neurol Genet, 6, e425.

29. Yeganeh, M. and Hernandez, N. (2020) RNA polymerase III transcription as a disease factor. Genes Dev, 34, 865–882.

30. Merheb, E., Cui, M.H., DuBois, J.C., Branch, C.A., Gulinello, M., Shafit-Zagardo, B., Moir, R.D. and Willis, I.M. (2021) Defective myelination in an RNA polymerase III mutant leukodystrophic mouse. Proc Natl Acad Sci U S A, 118.

31. Michell-Robinson, M.A., Watt, K.E.N., Grouza, V., Macintosh, J., Pinard, M., Tuznik, M., Chen, X., Darbelli, L., Wu, C.L., Perrier, S. et al. (2023) Hypomyelination, hypodontia and craniofacial abnormalities in a Polr3b mouse model of leukodystrophy. Brain, 146, 5070–5085.

32. Yeganeh, M., Praz, V., Cousin, P. and Hernandez, N. (2017) Transcriptional interference by RNA polymerase III affects expression of the Polr3e gene. Genes Dev, 31, 413–421.

33. Cho, H., Lee, W., Kim, G.W., Lee, S.H., Moon, J.S., Kim, M., Kim, H.S. and Oh, J.W. (2019) Regulation of La/SSB-dependent viral gene expression by pre-tRNA 3’ trailer-derived tRNA fragments. Nucleic Acids Res, 47, 9888–9901.

34. Gahl, W.A., Markello, T.C., Toro, C., Fajardo, K.F., Sincan, M., Gill, F., Carlson-Donohoe, H., Gropman, A., Pierson, T.M., Golas, G. et al. (2012) The National Institutes of Health Undiagnosed Diseases Program: insights into rare diseases. Genet Med, 14, 51–59.

35. Gahl, W.A., Mulvihill, J.J., Toro, C., Markello, T.C., Wise, A.L., Ramoni, R.B., Adams, D.R., Tifft, C.J. and Udn. (2016) The NIH Undiagnosed Diseases Program and Network: Applications to modern medicine. Mol Genet Metab, 117, 393–400.

36. Richards, S., Aziz, N., Bale, S., Bick, D., Das, S., Gastier-Foster, J., Grody, W.W., Hegde, M., Lyon, E., Spector, E. et al. (2015) Standards and guidelines for the interpretation of sequence variants: a joint consensus recommendation of the American College of Medical Genetics and Genomics and the Association for Molecular Pathology. Genet Med, 17, 405–424.

37. Robinson, A., Escuin, S., Doudney, K., Vekemans, M., Stevenson, R.E., Greene, N.D., Copp, A.J. and Stanier, P. (2012) Mutations in the planar cell polarity genes CELSR1 and SCRIB are associated with the severe neural tube defect craniorachischisis. Hum Mutat, 33, 440–447.

38. Lei, Y., Zhu, H., Duhon, C., Yang, W., Ross, M.E., Shaw, G.M. and Finnell, R.H. (2013) Mutations in planar cell polarity gene SCRIB are associated with spina bifida. PLoS One, 8, e69262.

39. Kurosaki, T. and Maquat, L.E. (2016) Nonsense-mediated mRNA decay in humans at a glance. J Cell Sci, 129, 461–467.

40. Cormack, B.P. and Struhl, K. (1992) The TATA-binding protein is required for transcription by all three nuclear RNA polymerases in yeast cells. Cell, 69, 685–696.

41. Johnson, S.A., Dubeau, L. and Johnson, D.L. (2008) Enhanced RNA polymerase III-dependent transcription is required for oncogenic transformation. J Biol Chem, 283, 19184–19191.

42. Sethy-Coraci, I., Moir, R.D., Lopez-de-Leon, A. and Willis, I.M. (1998) A differential response of wild type and mutant promoters to TFIIIB70 overexpression in vivo and in vitro. Nucleic Acids Res, 26, 2344–2352.

43. Romanelli, V., Nakabayashi, K., Vizoso, M., Moran, S., Iglesias-Platas, I., Sugahara, N., Simon, C., Hata, K., Esteller, M., Court, F. et al. (2014) Variable maternal methylation overlapping the nc886/vtRNA2-1 locus is locked between hypermethylated repeats and is frequently altered in cancer. Epigenetics, 9, 783–790.

44. Silver, M.J., Kessler, N.J., Hennig, B.J., Dominguez-Salas, P., Laritsky, E., Baker, M.S., Coarfa, C., Hernandez-Vargas, H., Castelino, J.M., Routledge, M.N. et al. (2015) Independent genomewide screens identify the tumor suppressor VTRNA2-1 as a human epiallele responsive to periconceptional environment. Genome Biol, 16, 118.

45. Park, J.L., Lee, Y.S., Song, M.J., Hong, S.H., Ahn, J.H., Seo, E.H., Shin, S.P., Lee, S.J., Johnson, B.H., Stampfer, M.R. et al. (2017) Epigenetic regulation of RNA polymerase III transcription in early breast tumorigenesis. Oncogene, 36, 6793–6804.

46. Lee, Y.S. (2022) Are We Studying Non-Coding RNAs Correctly? Lessons from nc886. Int J Mol Sci, 23.

47. Rijal, K. and Maraia, R.J. (2013) RNA polymerase III mutants in TFIIFα-like C37 cause terminator readthrough with no decrease in transcription output. Nucleic Acids Research, 41, 139–155.

48. Mishra, S., Hasan, S.H., Sakhawala, R.M., Chaudhry, S. and Maraia, R.J. (2021) Mechanism of RNA Polymerase III termination-associated reinitiation-recycling conferred by the essential function of the N terminal-and-Linker domain of the C11 subunit. Nat Commun, 12, 5900.

49. Canella, D., Bernasconi, D., Gilardi, F., Lemartelot, G., Migliavacca, E., Praz, V., Cousin, P., Delorenzi, M. and Hernandez, N. (2012) A multiplicity of factors contributes to selective RNA polymerase III occupancy of a subset of RNA polymerase III genes in mouse liver. Genome Res, 22, 666–680.

50. Thornlow, B.P., Armstrong, J., Holmes, A.D., Howard, J.M., Corbett-Detig, R.B. and Lowe, T.M. (2020) Predicting transfer RNA gene activity from sequence and genome context. Genome Res, 30, 85–94.

51. Goodier, J.L. and Maraia, R.J. (1998) Terminator-specific recycling of a B1-Alu transcription complex by RNA polymerase III is mediated by the RNA terminus-binding protein La. J Biol Chem, 273, 26110–26116.

52. Yoo, C.J. and Wolin, S.L. (1997) The yeast La protein is required for the 3’ endonucleolytic cleavage that matures tRNA precursors. Cell, 89, 393–402.

53. Intine, R.V.A., Sakulich, A.L., Koduru, S.B., Huang, Y., Pierstorrf, E., Goodier, J.L., Phan, L. and Maraia, R.J. (2000) Control of transfer RNA maturation by phosphorylation of the human La antigen on serine 366. Mol Cell, 6, 339–348.

54. Wolin, S.L. and Cedervall, T. (2002) The La protein. Annu Rev Biochem, 71, 375–403.

55. Maraia, R.J., Chang, D.Y., Wolffe, A.P., Vorce, R.L. and Hsu, K. (1992) The RNA polymerase III terminator used by a B1-Alu element can modulate 3’ processing of the intermediate RNA product. Mol Cell Biol, 12, 1500–1506.

56. Maraia, R.J., Kenan, D.J. and Keene, J.D. (1994) Eukaryotic transcription termination factor La mediates transcript release and facilitates reinitiation by RNA polymerase III. Mol. Cell. Biol., 14, 2147–2158.

57. Pannone, B., Xue, D. and Wolin, S.L. (1998) A role for the yeast La protein in U6 snRNP assembly: evidence that the La protein is a molecular chaperone for RNA polymerase III transcripts. EMBO J, 17, 7442–7453.

58. Lee, K., Kunkeaw, N., Jeon, S.H., Lee, I., Johnson, B.H., Kang, G.Y., Bang, J.Y., Park, H.S., Leelayuwat, C. and Lee, Y.S. (2011) Precursor miR-886, a novel noncoding RNA repressed in cancer, associates with PKR and modulates its activity. RNA, 17, 1076–1089.

59. Kickhoefer, V.A., Poderycki, M.J., Chan, E.K. and Rome, L.H. (2002) The La RNA-binding protein interacts with the vault RNA and is a vault-associated protein. J Biol Chem, 277, 41282–41286.

60. Parrott, A.M. and Mathews, M.B. (2007) Novel rapidly evolving hominid RNAs bind nuclear factor 90 and display tissue-restricted distribution. Nucleic Acids Res, 35, 6249–6258.

61. Younis, I., Dittmar, K., Wang, W., Foley, S.W., Berg, M.G., Hu, K.Y., Wei, Z., Wan, L. and Dreyfuss, G. (2013) Minor introns are embedded molecular switches regulated by highly unstable U6atac snRNA. Elife, 2, e00780.

62. Belair, C., Sim, S. and Wolin, S.L. (2018) Noncoding RNA Surveillance: The Ends Justify the Means. Chem Rev, 118, 4422–4447.

63. Porat, J., Kothe, U. and Bayfield, M.A. (2021) Revisiting tRNA chaperones: New players in an ancient game. RNA, 27, 543–559.

64. Gerber, A., Ito, K., Chu, C.S. and Roeder, R.G. (2020) Gene-Specific Control of tRNA Expression by RNA Polymerase II. Mol Cell, 78, 765–778.e767.

65. Jiang, Y., Huang, J., Tian, K., Yi, X., Zheng, H., Zhu, Y., Guo, T. and Ji, X. (2022) Cross-regulome profiling of RNA polymerases highlights the regulatory role of polymerase III on mRNA transcription by maintaining local chromatin architecture. Genome Biol, 23, 246.

66. Kumar, P., Mudunuri, S.B., Anaya, J. and Dutta, A. (2015) tRFdb: a database for transfer RNA fragments. Nucleic Acids Res, 43, D141–145.

67. Lee, Y.S., Shibata, Y., Malhotra, A. and Dutta, A. (2009) A novel class of small RNAs: tRNA-derived RNA fragments (tRFs). Genes Dev, 23, 2639–2649.

68. Wilson, B., Su, Z., Kumar, P. and Dutta, A. (2023) XRN2 suppresses aberrant entry of tRNA trailers into argonaute in humans and Arabidopsis. PLoS Genet, 19, e1010755.

69. Yang, A., Bofill-De Ros, X., Stanton, R., Shao, T.J., Villanueva, P. and Gu, S. (2022) TENT2, TUT4, and TUT7 selectively regulate miRNA sequence and abundance. Nat Commun, 13, 5260.

70. Hasler, D., Lehmann, G., Murakawa, Y., Klironomos, F., Jakob, L., Grasser, F.A., Rajewsky, N., Landthaler, M. and Meister, G. (2016) The Lupus Autoantigen La Prevents Mis-channeling of tRNA Fragments into the Human MicroRNA Pathway. Mol Cell, 63, 110–124.

71. Wang, Z., Bai, L., Hsieh, Y. and Roeder, R.G. (2000) Nuclear factor 1 (NF1) affects accurate termination and multiple-round transcription by human RNA polymerase III. EMBO J, 19, 6823–6832.

72. Wang, Z. and Roeder, R.G. (1998) DNA topoisomerase I and PC4 can interact with human TFIIIC to promote both accurate termination and transcription reinitiation by RNA polymerase III. Mol. Cell, 1, 749–757.

73. Gottlieb, E. and Steitz, J.A. (1989) Function of the mammalian La protein: evidence for its action in transcription termination by RNA polymerase III. EMBO J., 8, 851–861.

74. Goodier, J.L., Fan, H. and Maraia, R.J. (1997) A carboxy-terminal basic region controls RNA polymerase III transcription factor activity of human La protein. Mol Cell Biol, 17, 5823–5832.

75. Weser, S., Bachmann, M., Seifart, K.H. and Meißner, W. (2000) Transcription efficiency of human polymerase III genes in vitro does not depend on the RNP-forming autoantigen La. Nucl. Acids. Res., 28, 3935–3942.

76. Dieci, G., Bosio, M.C., Fermi, B. and Ferrari, R. (2013) Transcription reinitiation by RNA polymerase III. Biochim Biophys Acta, 1829, 331–341.

77. Stefano, J.E. (1984) Purified lupus antigen La recognizes an oligouridylate stretch common to the 3’ termini of RNA polymerase III transcripts. Cell, 36, 145–154.

78. Nashimoto, M., Nashimoto, C., Tamura, M., Kaspar, R.L. and Ochi, K. (2001) The inhibitory effect of the autoantigen La on in vitro 3’ processing of mammalian precursor tRNAs. J Mol Biol, 312, 975–984.

79. Liang, C., Xiong, K., Szulwach, K.E., Zhang, Y., Wang, Z., Peng, J., Fu, M., Jin, P., Suzuki, H.I. and Liu, Q. (2013) Sjogren syndrome antigen B (SSB)/La promotes global microRNA expression by binding microRNA precursors through stem-loop recognition. J Biol Chem, 288, 723–736.

80. Zheng, Q., Yang, H.J. and Yuan, Y.A. (2017) Autoantigen La Regulates MicroRNA Processing from Stem-Loop Precursors by Association with DGCR8. Biochemistry, 56, 6098–6110.

81. Stentenbach, M., Ermer, J.A., Rudler, D.L., Perks, K.L., Raven, S.A., Lee, R.G., McCubbin, T., Marcellin, E., Siira, S.J., Rackham, O. et al. (2023) Multi-omic profiling reveals an RNA processing rheostat that predisposes to prostate cancer. EMBO Mol Med, 15, e17463.

82. Siira, S.J., Rossetti, G., Richman, T.R., Perks, K., Ermer, J.A., Kuznetsova, I., Hughes, L., Shearwood, A.J., Viola, H.M., Hool, L.C. et al. (2018) Concerted regulation of mitochondrial and nuclear non-coding RNAs by a dual-targeted RNase Z. EMBO Rep, 19.

83. Bayfield, M.A. and Maraia, R.J. (2009) Precursor-product discrimination by La protein during tRNA metabolism. Nat Struct & Mol Biol, 16, 430–437.

84. Sommer, G. and Heise, T. (2020) Role of the RNA-binding protein La in cancer pathobiology. RNA Biol, 1–19.

85. Stephen, J., Maddirevula, S., Nampoothiri, S., Burke, J.D., Herzog, M., Shukla, A., Steindl, K., Eskin, A., Patil, S.J., Joset, P. et al. (2018) Bi-allelic TMEM94 Truncating Variants Are Associated with Neurodevelopmental Delay, Congenital Heart Defects, and Distinct Facial Dysmorphism. Am J Hum Genet, 103, 948–967.

86. Fairley, J.A., Kantidakis, T., Kenneth, N.S., Intine, R.V., Maraia, R.J. and White, R.J. (2005) Human La is Found at RNA Polymerase III-Transcribed Genes In Vivo. Proc Nat Acad Sci, USA, 102, 18350–18355.

87. Maraia, R. (1991) The subset of mouse B1 (*Alu*-equivalent) sequences expressed as small processed cytoplasmic transcripts. Nucl. Acids Res., 19, 5695–5702.

88. Leonard G. Davis, M.D.D.a.J.F.B. (1986) Basic Methods in Molecular Biology. Elsevier.

89. Dobin, A., Davis, C.A., Schlesinger, F., Drenkow, J., Zaleski, C., Jha, S., Batut, P., Chaisson, M. and Gingeras, T.R. (2013) STAR: ultrafast universal RNA-seq aligner. Bioinformatics, 29, 15–21.

90. Liao, Y., Smyth, G.K. and Shi, W. (2014) featureCounts: an efficient general purpose program for assigning sequence reads to genomic features. Bioinformatics, 30, 923–930.

91. Anders, S. and Huber, W. (2010) Differential expression analysis for sequence count data. Genome Biol, 11, R106.

92. Love, M.I., Huber, W. and Anders, S. (2014) Moderated estimation of fold change and dispersion for RNA-seq data with DESeq2. Genome Biol, 15, 550.

93. Wu, T., Hu, E., Xu, S., Chen, M., Guo, P., Dai, Z., Feng, T., Zhou, L., Tang, W., Zhan, L. et al. (2021) clusterProfiler 4.0: A universal enrichment tool for interpreting omics data. Innovation (N Y), 2, 100141.

## REFERENCES for Supplementary Materials

1. Robinson, A. et al. Mutations in the planar cell polarity genes CELSR1 and SCRIB are associated with the severe neural tube defect craniorachischisis. Hum Mutat 33, 440–7 (2012).

2. How, J.Y., Stephens, R., Lim, K.Y.B., Humbert, P.O. & Kvansakul, M. Structural basis of the human Scribble-Vangl2 association in health and disease. Biochem J BCJ20200816. (2021).

3. Zarbalis, K. et al. A focused and efficient genetic screening strategy in the mouse: identification of mutations that disrupt cortical development. PLoS Biol 2, E219 (2004).

4. Arimbasseri, A.G. et al. RNA polymerase III output is functionally linked to tRNA dimethyl-G26 modification. PLoS Genetics 11, e1005671 (2015).

5. Rijal, K., Maraia, RJ & Arimbasseri, AG. A methods review on use of nonsense suppression to study 3’ end formation and other aspects of tRNA biogenesis. Gene 556, 35–50 (2015).

6. Iben, J.R. et al. Point mutations in the Rpb9-homologous domain of Rpc11 that impair transcription termination by RNA polymerase III. Nucleic Acids Res 39, 6100–6113 (2011).

7. Rijal, K. & Maraia, R.J. Active Center Control of Termination by RNA Polymerase III and tRNA Gene Transcription Levels In Vivo. PLoS Genet 12, e1006253 (2016).

8. Mishra, S., Hasan, S.H., Sakhawala, R.M., Chaudhry, S. & Maraia, R.J. Mechanism of RNA Polymerase III termination-associated reinitiation-recycling conferred by the essential function of the N terminal-and-Linker domain of the C11 subunit. Nat Commun 12, 5900 (2021).

9. Arimbasseri, A.G., Kassavetis, G.A. & Maraia, R.J. Comment on “Mechanism of eukaryotic RNA polymerase III transcription termination”. Science 345, 524 (2014).

10. Perrier, S. et al. Expanding the phenotypic and molecular spectrum of RNA polymerase III-related leukodystrophy. Neurol Genet 6, e425 (2020).

11. Wolf, N.I. et al. Clinical spectrum of 4H leukodystrophy caused by POLR3A and POLR3B mutations. Neurology 83, 1898–905 (2014).

12. de Assis Pereira Matos, P.C.A., et al. POLR3A-Related Disorder Presenting with Late-Onset Dystonia and Spastic Paraplegia. Mov Disord Clin Pract 7, 467–469 (2020).

13. Kashiki, H. et al. POLR1C variants dysregulate splicing and cause hypomyelinating leukodystrophy. Neurol Genet 6, e524 (2020).

14. Báez-Becerra, C.T. et al. Nucleolar disruption, activation of P53 and premature senescence in POLR3A-mutated Wiedemann-Rautenstrauch syndrome fibroblasts. Mech Ageing Dev 192, 111360 (2020).

15. Beauregard-Lacroix, E. et al. A variant of neonatal progeroid syndrome, or Wiedemann-Rautenstrauch syndrome, is associated with a nonsense variant in POLR3GL. Eur J Hum Genet (2020).

16. Jay, A.M. et al. Neonatal progeriod syndrome associated with biallelic truncating variants in POLR3A. Am J Med Genet A 170, 3343–3346 (2016).

17. Paolacci, S. et al. Specific combinations of biallelic POLR3A variants cause Wiedemann-Rautenstrauch syndrome. J Med Genet 55, 837–846 (2018).

18. Wambach, J.A. et al. Bi-allelic POLR3A Loss-of-Function Variants Cause Autosomal-Recessive Wiedemann-Rautenstrauch Syndrome. Am J Hum Genet 103, 968–975 (2018).

19. Kulhánek, J. et al. POLR3B-associated leukodystrophy: clinical, neuroimaging and molecular-genetic analyses in four patients: clinical heterogeneity and novel mutations in POLR3B gene. Neurol Neurochir Pol 53, 369–376 (2019).

20. Gauquelin, L. et al. Clinical spectrum of POLR3-related leukodystrophy caused by biallelic POLR1C pathogenic variants. Neurol Genet 5, e369 (2019).

21. Richards, S. et al. Standards and guidelines for the interpretation of sequence variants: a joint consensus recommendation of the American College of Medical Genetics and Genomics and the Association for Molecular Pathology. Genet Med 17, 405–24 (2015).

22. Stephen, J. et al. Cellular and molecular defects in a patient with Hermansky-Pudlak syndrome type 5. PLoS One 12, e0173682 (2017).

